# Dispensable genome and segmental duplications drive the genome plasticity in *Fusarium solani*

**DOI:** 10.1101/2024.06.05.597655

**Authors:** Abbeah Navasca, Jatinder Singh, Viviana Rivera-Varas, Upinder Gill, Gary Secor, Thomas Baldwin

**Affiliations:** Department of Plant Pathology, North Dakota State University, Fargo, ND, USA

**Keywords:** *Fusarium solani*, pan-genome, accessory chromosomes, segmental duplications, genome plasticity

## Abstract

*Fusarium solani* is a species complex encompassing a large phylogenetic clade with diverse members occupying varied habitats. We recently reported a unique opportunistic *F. solani* associated with unusual dark galls in sugarbeet. We assembled the chromosome-level genome of the *F. solani* sugarbeet isolate strain SB1 using Oxford Nanopore and Hi-C sequencing. SB1 has a large genome (59.38 Mb) organized into 15 chromosomes. The genome expansion is due to the high repeats and massive segmental duplications within its three potentially accessory chromosomes. These chromosomes are absent in the closest reference genome with chromosome-level assembly, *F. vanettenii* 77-13-4. The extensive segmental duplications between the two SB1 chromosomes suggest that this isolate may have doubled its accessory genes. Further comparison of the *F. solani* strain SB1 genome demonstrates inversions and syntenic regions to an accessory chromosome of *F. vanettenii* 77-13-4. The pan-genome of 12 publicly available *F. solani* isolates nearly reached gene saturation, with few new genes discovered after the addition of the last genome. Based on orthogroups and average nucleotide identity, *F. solani* is not grouped by lifestyle or origin. The pan-genome analysis further revealed the enrichment of several enzymes-coding genes within the dispensable (accessory + unique genes) genome, such as hydrolases, transferases, oxidoreductases, lyases, ligases, isomerase, and dehydrogenase. The evidence presented here suggests that genome plasticity, genetic diversity, and adaptive traits in *Fusarium solani* are driven by the dispensable genome with significant contributions from segmental duplications.

## 1 Introduction

The *Fusarium solani* species complex (FSSC) comprises at least 77 phylogenetically distinct species known to thrive in diverse environments covering ecological, agricultural, and clinical settings (O’Donnell et al., 2020; Geiser et al., 2021). The FSSC has three distinct clades, denoted as Clades 1 through 3. Clade 1 comprises *F. plagianthi* and *F. illudens*, while Clade 2 contains several *Fusarium* species, including those that caused sudden death syndrome and bean root rot. Clade 3 is the largest, boasting over 60 distinct species. This clade houses clinically and agriculturally important species, including *Fusarium solani* (O’Donnell et al., 2008; O’Donnell et al., 2020; Geiser et al., 2021).

The trans-kingdom fungus *F. solani* is a highly adaptive species. In humans, *F. solani* affects immunocompromised patients causing skin and nail infections (Gupta et al., 2000; Godoy et al., 2004; Zhang et al., 2006; Kuruvilla et al., 2012), mycotic keratitis (Ahearn et al., 2008; Boral et al., 2018; Toravato et al., 2021) even in a healthy individual (Ortega-Rosales et al., 2019), and chronic diabetic ulcers (Pai et al., 2010). A recent case involving *F. solani* is the clinical meningitis outbreak in Mexico affecting Mexican and US residents who traveled for medical purposes (Smith et al., 2023; García-Rodríguez et al., 2024; Strong et al., 2024). *F. solani* is also a threat to animals. This fungus caused keratitis in rabbits (Kiryu et al., 1991), cutaneous hyalohyphomycosis and mass mortalities in loggerhead sea turtles (Cabanes et al., 1997; Sarmiento-Ramírez et al., 2010), tissue destruction and inflammation in shrimps and prawns (Hose et al. 1984; Bian et al., 1981; Le et al., 2005), dermatitis in false killer whales (Tanaka et al., 2012), and submucosal nodules on the face, tongue, and waist of a Doberman dog (Kano et al., 2001). Undoubtedly, *F. solani* infects crops across various species. Several reports highlight this pathogen’s ability to cause multiple rot diseases such as dry rot in potato stems (Goss, 1940), fruit rot in pumpkin (Rampersad, 2008), sweet pepper (Ramdial and Rampersad, 2010), and strawberry (Mehmood et al., 2017); crown rot in cucumber (Li et al., 2010) and strawberry (Pastrana et al., 2014; Villarino et al., 2019); root rot in peas (Van Etten, 1978; Gibert et al., 2023), sweet potato (Wang et al., 2013), strawberry (Pastrana et al., 2014; Villarino et al., 2019), okra (Lie et al., 2015), eggplant (Li et al., 2017), tobacco (Yang et al., 2020), and many more. Apart from fruit and vegetables, *F. solani* also infects ornamental plants such as bulb rot in tulips (Nisa et al., 2021), soft rot (Han et al., 2017) and wilt (Xie et al., 2024) in orchids. Other symptoms caused by *F. solani* are cankers in sweet potato (Wang et al., 2013) and English Walnut (Chen and Swart, 2007; Mulero-Aparicio et al., 2019; Tuerdi et al., 2023), gummosis in rubber trees (Huang et al., 2016), wilt in cotton (Zhu et al., 2019), leaf-sheath rot in bush lily (Sun et al., 2022), and leaf spot in pineapple (Liao et al., 2024).

Aside from being an important pathogen of humans, animals, and plants, *F. solani* also have other lifestyles. It thrives as an endophyte in mulberry (Kim et al., 2017) and arabidopsis (Mesny et al., 2021) and also as a saprophyte, increasing the insecticidal efficacy of the entomopathogenic nematode *Steinernema diaprepesi* (Wu et al., 2018). Recently, we reported *F. solani* as an opportunistic pathogen of sugarbeet (Navasca et al., 2023). This isolate, *F. solani* strain SB1, was recovered from dark galls of sugarbeets, with symptoms that differ from other *Fusarium* diseases in sugarbeets (Hanson, 2006; Rivera et al., 2008; Secor et al., 2014; Khan et al., 2021). Electron microscopy and sequencing work confirmed that the galls contain sugarbeet material. However, greenhouse tests show that SB1 can only cause mild vascular discoloration without developing any galls or gall-like structures, adding a layer of complexity to the disease.

The broad host range and adaptive lifestyle of *Fusarium* species can be attributed to their ability to take in genomic regions, making them very adaptive to specific environments (Coleman et al., 2009; Ma et al., 2010; Ma et al., 2013). Knowing this ability, we sequenced the *F. solani* strain SB1 to determine its genetic elements and how it compares to *F. solani* genomes, which could provide insight into this unique species and disease complex. The concept of supernumerary chromosomes, more commonly called accessory chromosomes (ACs), is well-established in *Fusarium*, particularly *Fusarium solani* and *Fusarium oxysporum* (Coleman et al., 2009; Ma et al., 2010; Ma et al., 2013; Zhang et al., 2020). These ACs are not essential for growth but offer added features such as pathogenicity or increased virulence. We hypothesize that *F. solani* SB1 contains accessory regions that enable it to be opportunistic to sugarbeet and are unique from other *F. solani* species. Here, we describe its complete genomic characteristics and how it compares with the genome of the pea pathogen, *Fusarium vanettenii* MPVI isolate 77-13-4 (previously reported as *N. haematococca*; Coleman et al., 2009). Given the ability of *F. solani* to thrive in various habitats, relying on reference genomes may overlook crucial adaptive genes present in unrepresented isolates. In this study, we performed a pan-genome analysis of publicly available twelve *F. solani* genomes encompassing pathogens of plants and animals, saprophytes, and endophytes, including the opportunistic pathogen of sugarbeet. We determined the enrichment of core and dispensable genes. Our findings underscore the remarkable genome plasticity, genetic diversity, and inherent adaptive ability of *Fusarium solani*.

## 2 Methodology

### 2.1 Isolation, DNA Extraction, and Draft Genome Sequencing

Procedures for isolation, DNA extraction, library preparation, and draft genome sequencing are available in Navasca et al., 2023. In brief, we isolated *Fusarium solani* from galled sugarbeet by excising and sterilizing small tissues with 0.5% v/v sodium hypochlorite for 10 minutes and washing with sterile distilled water. We extracted high-molecular-weight DNA from the mycelia of pure culture *F. solani* sugarbeet isolate strain SB1 grown in PDB for five days following the instructions of Liu et al. (2021). Agilent TapeStation and Qubit 4.0 determined the DNA quality and quantity, respectively. We utilized the Nanopore Protocol Lambda Control Experiment (SQK-LSK109) for library preparation and performed sequencing using the R10.3 version flowcell in MinIOn. Guppy version 6.0.1 (Oxford Nanopore Technologies, UK) basecalled the reads, followed by adapter removal using Porechop version 0.2.4 (Wick et al., 2017) and data quality control using LongQC version 1.2.0 (Fukasawa et al. 2021). Finally, we assembled the sequences using NECAT version 0.0.1 (Chen et al. 2021) and checked the quality using Quast 5.0.2 (Mikheenko et al. 2018).

### 2.2 Hi-C Sequencing and Chromosome-Level Genome Assembly

Mycelia of *F. solani* SB1 grown in PDA broth for five days were collected and resuspended in 1% formaldehyde in molecular-grade water, followed by a 20-minute incubation with periodic mixing. Glycine (125 mM final concentration) was then added to the sample and incubated for 15 min with occasional mixing. A final spin down (1000 *g*) for 5 min separated the sample from the mixture, and the supernatant was removed before sending the sample to Phase Genomics for ProxiMeta high-throughput chromosome conformation capture or Hi-C sequencing on an Illumina NovaSeq system (Phase Genomics, Seattle, Washington, USA). Genome completeness was assessed separately by Fungal Genome Mapping Project (FGMP) version 1.0.2 (Cissé and Stajich, 2019) and Benchmarking Universal Single-Copy Orthologs (BUSCO) version 5.2.2 (Simão et al., 2015; Manni et al., 2021). In BUSCO analysis, *F. graminearum* gene model from Augustus version 3.4.0 (Stanke and Waack, 2003) was used for evaluating the genome completeness. Telomeric regions were identified by the ‘TTAGGG’ sequence specific for most ascomycetes (https://telomerase.asu.edu).

### 2.3 Genome Annotation

We masked the genome and identified transposable elements via RepeatMasker version 4.0.9 (Smit et al., 2003) using a custom library produced from RepeatModeler2 version 2.0.5 (Flynn et al. 2020) with built-in three *de-novo* repeat finding programs RECON, RepeatScout, and LtrHarvest/Ltr_retriever. Evidence-based annotation was performed using Maker version 2.3.11 (Holt and Yandell, 2011) pipeline with built-in *ab-initio* gene annotation programs SNAP version 2013-11-29 (Korf, 2004) and Augustus version 3.4.0 (Stanke and Waack, 2003). We did three rounds of annotations utilizing ESTs and proteins of the well-studied reference genome *F. vanettenii* strain 77-13-4, formerly reported as *Nectria haematococca* mating population MPVI (Coleman et al., 2009), from EnsemblFungi (accessed November 12, 2023) and the reference genome *F. solani* strain FSSC 5 MPI-SDFR-AT-0091 (accessed November 28, 2023), reviewed proteins of *F. solani* from Uniprot (accessed December 14, 2023), and the custom library of repeats from RepeatModeler2 to produce a high-quality annotation for downstream analysis (Batut et al., 2018; Bretaudeau, 2023). We implemented the same annotation method to eight other *Fusarium solani* genomes from NCBI accessed on November 28, 2023 with only published raw data (Accessions: GCA_027574645.1, GCA_002215905.1, GCA_019320015.1, GCA_013168735.1, GCA_024220475.1, GCA_033085375.1, GCA_030014125.1, and GCA_029603225.1). Annotations of one genome from the submitter in NCBI (Accession: GCA_027945525.1) and *F. vanettenii* 77-13-4 v2.0, and *F. solani* FSSC 5 MPI-SDFR-AT-0091 (Accession: GCA_020744495.1) both from JGI, were used as is for further analysis. A total of 12 strains were used in this study. Following annotation, we determined genome completeness using proteins in BUSCO version 5.3.2 (Stanke and Waack, 2003; Manni et al., 2021) aided by *F. graminearum* genome. We estimated gene density over 100kb region of each chromosome of *F. solani* strain SB1.

### 2.4 Prediction of Pathogenicity-Related Genes

Genomic annotations were made for *F. solani* sugarbeet isolate strain SB1. Secondary metabolism potential was determined using the antiSMASH 7.0 fungal version (Blin et al. 2021) with the ‘relaxed’ option. SignalP 6.0 for eukaryotes (Teufel et al., 2022) and TargetP 2.0 with non-plant option (Almagro Armenteros et al., 2019) predicted the secretory signal peptides and mitochondrial proteins, respectively. After signal peptide prediction and removal of mitochondrial proteins, sequences were then subjected to DeepTMHMM 1.0.24 (Hallgren et al., 2022) and Phobius (Käll et al., 2004) to remove proteins with the transmembrane domain. The remaining protein sets were checked for glycosylphosphatidylinositol (GPI) anchors using NetGPI 1.1 (Gíslason et al., 2021). Ultimately, only proteins without GPI anchors were considered secreted proteins. The refined secreted proteins were further used for the analysis of pathogenesis-related proteins. The dbCAN3 (Zheng et al., 2023) meta server, which combines HMMER, DIAMOND, and eCAMI database, annotated the carbohydrate-active enzymes (CAZymes) while EffectorP 3.0 for fungi (Sperschneider J and Dodds, 2021) predicted the effector proteins. We performed a BLAST search in the Pathogen-Host Interaction Database or PHI-base version 4.0 with protein sequences 4.14 database (Urban et al., 2022) to find homologs of refined secreted proteins functionally characterized on other organisms. We used e-value <1e-05 and at least 50% identity to screen the hits. We selected the gene with the highest bit score for genes with more than one hit. Bit scores consider both the alignment of the sequences and the gaps within the alignment. A high bit score indicates a better alignment (Madden, 2003). Finally, we determined the gene function intersection between CAZymes, effectors, and PHIs.

### 2.5 Genome Comparisons and Gene Collinearity

We applied Mauve (Darling et al., 2004) plug-in software in Geneious Prime version 2023.1.2 to align the SB1 genome to itself to determine the global rearrangement structure within the SB1 genome. We utilized the alignment file to generate a plot using Circos version 0.69.8 (Krzywinski et al., 2009; Rasche and Hiltemann, 2020). We used Chromeister version 1.5a (Pérez-Wohlfeil et al., 2019) to fast-align DNA sequences and determine the sequence similarity of *F. solani* SB1 and *F. vanettenii* 77-13-4 v2.0. For synteny analysis, we performed all vs all protein alignment between SB1 and *F. vanettenii* using the BLASTP service of BLAST and prepared the bed files according to the requirements of McScanX (Wang et al., 2012). We also used the same procedure to identify gene collinearity within the SB1 genome with the duplicate_gene_classifier option in McScanX to determine the classification of gene duplications. Syntenic blocks were visualized in SynVisio (Bandi and Gutwin, 2020).

### 2.6 Pangenome Analysis and GO Enrichment

OrthoFinder version 2.5.5 (Emms and Kelly, 2019) identified clusters of orthologous proteins between 12 strains. The pangenome of *F. solani* was analyzed using core and dispensable proteins and was visualized in a curve in PanGP (Zhao et al., 2014; Li et al., 2022) in ‘totally random’ algorithm at 1000 combinations, replicated 50 times. Core proteins are those consistently found across all strains, whereas dispensable proteins cover accessory proteins (present in two or more strains) and unique proteins (exclusive to a single strain). Functional annotation of core and dispensable (accessory and unique) proteins was performed using InterProScan version 5.59-91.0 (Jones et al., 2014). We utilized ShinyGo version 0.80 (Ge et al., 2020) with false discovery rate (FDR) correction at p < 0.05 to evaluate the enrichment of core and dispensable genomes against the *F. vanetteni* genome.

### 2.7 Phylogenomic Analysis

Protein sequences of *F. solani* strains in this study and those of the two outgroups *Fusarium graminearum* PH-1 NNRL 31084 (Accession: GCA_000240135.3) and *Fusarium oxysporum* f. sp. *lycopersici* 4287 strain (Accession: GCA_000149955.2) were utilized for phylogenomic analysis using OrthoFinder version 2.5.5 (Emms and Kelly 2017; Emms and Kelly, 2019). We used the software’s default parameters to produce the species tree using the information of the entire set of orthologous groups present in all *F. solani* strains. The tree was visualized using iTol version 6.8.1 (Letunic and Bork, 2021). We also computed for the average nucleotide identity of *F. solani* strains using FastANI version 1.3 (Jain et al., 2018).

## 3.0 Results

### 3.1 Genome Features of *Fusarium solani* strain SB1

The draft genome assembly of *F. solani* SB1 was assembled previously in 19 contigs (Navasca et. al., 2023) using Oxford Nanopore sequencing. High-throughput chromatin conformation capture (Hi-C) sequencing resolved the genome assembly in 15 chromosomes with a total size of 59.4 Mb and N50 of 4,122,546 bp with the largest contig at 6,537,432 bp. The chromosome size ranges from 6.54 to 2.11 Mb (Figure 1A). We achieved 96.1% genome completeness in Fungal Genome Mapping Project (FGMP) version 1.0.2 (Cissé and Stajich, 2019) and 98.5% in BUSCO version 5.2.2 (Simão et al., 2015; Manni et al., 2021). We found 13 chromosomes containing telomeric repeats at one end (Figure 1B). GC content of chromosomes ranged from 47 to 53% (Figure 1C). We also assembled the mitochondrial genome of *F. solani* SB1 isolate. The SB1 genome contains 17,981 protein-coding genes predicted by MAKER2 version 2.3.11 (Holt and Yandell, 2011) and 41 secondary metabolites identified by antiSMASH 7.0 fungal version (Blin et al. 2021) and 10.71% repetitive elements. Screening of protein-coding genes by SignalP 6.0 for eukaryotes (Teufel et al., 2022) and TargetP 2.0 (Almagro Armenteros et al., 2019), DeepTMHMM version 1.0.24 (Hallgren et al., 2022), Phobius (Käll et al., 2004), and NetGPI (Gislason et al., 2021) predicted 1,177 secreted proteins. EffectorP 3.0 for fungi (Sperschneider J and Dodds, 2021) recognized a total of 440 putative effector proteins (37% of secreted proteins) classified as either cytoplasmic (194) or apoplastic (246) where at least 135 are potentially dual-localized effectors (cytoplasmic/apoplastic or apoplastic/cytoplasmic). A total of 308 carbohydrate-active enzymes (26% of secreted proteins) were predicted by the dbCAN3 meta server (Zhang et al., 2018). We found 141 proteins (12% of secreted proteins) having potential roles in pathogenesis from the BLAST results of PHI-base version 4 (Urban et al., 2022) based on the parameters we set (homology > 50%, e-value <1e-05). All data are available in Supplementary Table 1.

**Figure 1.**
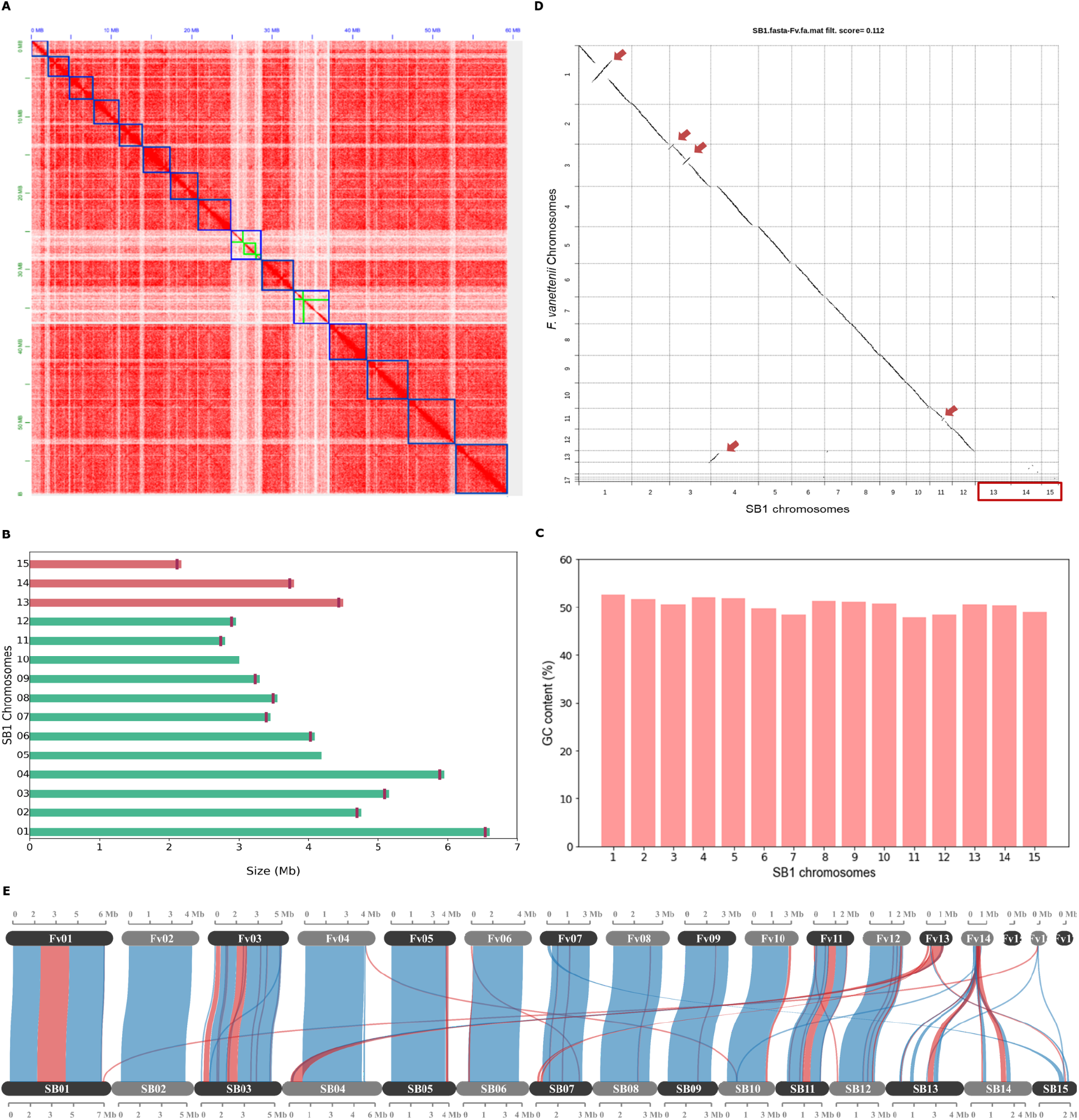
The *F. solani* SB1 genome assembly and comparison with *F. vanettenii* strain 77-13-4. **(A)** Hi-C contact map of SB1 scaffolds, outlined with blue squares, correspond to the 15 chromosomes sorted according to their length. The color intensity represents the frequency of contact between two chromosomes; **(B)** The *F. solani* SB1 chromosomes arranged according to similarity to the reference genome *F. vanettenii* strain 77-13-4. Three chromosomes, Chr13, Chr14, and Chr15 (pink bars) are not found in the reference genome. Note the vertical lines marking the telomeres of each chromosome except Chr05 and Chr10; **(C)** GC content is similar across SB1 chromosomes; **(D)** Genome comparison of *F. solani* SB1 and *F. vanettenii* shows high sequence similarity with inverted regions indicated by red arrows. Three chromosomes (Chr13, Chr14, and Chr15) are not found in *F. vanettenii*, indicated by the red box; **(E)** Gene collinearity *F. vanettenii* 77-13-4 and *F. solani* SB1 shows syntenic (blue) and inverted regions (red). SB1 Chr13 and Chr14 have collinear genes with the accessory chromosome Chr14 of *F. vanettenii*. Fv - *F. vanetenii* 77-13-4, SB - *F. solani* SB1.

### 3.2 *F. solani* SB1 harbors chromosomes not found in *F. vanettenii* strain 77-13-4

*Fusarium vanettenii* strain 77-13-4 was previously named *Nectria haematococca* (Coleman et al., 2009) and is the most studied strain in *Fusarium solani* group (Miao et al., 1991; Kistler et al., 1996; Wasmann and VanEtten, 1996; Han et al., 2001; Liu et al., 2003; Coleman et al., 2009). For this reason, we chose to compare the *F. solani* strain SB1 to this reference genome. We reversed-complemented five scaffolds (2, 3, 7, 10, 11) of the *F. solani* SB1 isolate and named them according to their matches with the chromosomes of the *F. vanettenii* strain 77-13-4 (Figures 1A and 1D). Direct pairwise genome comparison of *F. solani* SB1 and *F. vanettenii* 77-13-4 in Chromeister version 1.5a (Pérez-Wohlfeil et al., 2019) obtained a 0.112 score (‘zero’ as the perfect score for similarity), which indicates genomes are very similar but they also contain inversions indicated by red arrows in Figure 1D. Moreover, three chromosomes, Chr13, Chr14, and Chr15, are predominantly absent from *F. vanettenii*, highlighted by the red box in Figure 1D. These chromosomes will be highlighted in the succeeding sections. Further analysis using protein-coding genes in McScanX (Wang et al., 2012) visualized in SynVisio (Bandi and Gutwin, 2020) shows that *F. solani* SB1 and *F. vanettenii* 77-13-4 share 79.16% collinear genes but also harbor inversions indicated by red links in Figure 1E. Using our raw reads, we confirmed the inversions in our assembly by manually checking and ensuring that contigs on those areas overlap and do not have gaps. *F. vanettenii* harbor three ACs, Chr14, Chr15, and Chr17 (Coleman et al., 2009). We found collinear regions in Chr13 and Chr14 of *F. solani* strain SB1 with the AC Chr14 of *F. vanettenii* 77-13-4 (Figure 1E), suggesting these SB1 chromosomes could also be accessory. We checked these collinear genes and found that most are uncharacterized proteins of *F. vanettenii* (data not shown).

### 3.3 Genomic Attributes of Chr13, Chr14, and Chr15

Among SB1 chromosomes, Chr13 had the highest repetitive elements, which make up a quarter of its genome at 26.34%, followed by Chr14 at 21.00% and Chr15 at 9.66% (Figure 2A). Class II DNA transposons mainly occupy these chromosomes, with 11% for Chr13 and Chr14 and about 4% for Chr15. Chr06 also contains Class II DNA transposons, while Chr06, Chr07, and Chr11 all contain Class I Retrotransposons. The average gene density of *F. solani* SB1 genome is 303 genes per Mb, but Chr13, Chr14, and Chr15 only have 206, 193, and 243 genes per Mb, respectively (Supplementary Table 1; Figure 2B). This is likely due to the highly repetitive elements in these chromosomes. Each chromosome has at least one predicted biosynthetic gene cluster (BCG) except Chr09, Chr13, and Chr14. These BCGs are classified as terpenes (3), phosphonate (1), isocyanide (1), fungal-RiPP-like (2), non-ribosomal peptide synthase, NRPS (18), and polyketide synthase, PKS (13), and NRPS-PKS hybrid (3). Three of the BCGs have 100% similarity to sansalvamide, choline, and ochratoxin A; three with 84-85% similarity to fusarubin, lucilactaean, and cyclosporin, one with 62% similarity with matachelin, while four have 28-40% similarity to squalestatin S1, gibepyrone-A, oxyjavanicin, and duclauxin. The remaining BCGs do not have similarities with known clusters. Among these known BCGs, sansalvamide, a cytotoxic cyclic depsipeptide, and fusarubin, a polyketide, are secondary metabolites produced by marine *Fusarium* and *F. fujikoroi*, respectively (Belofsky et al., 1999; Studt et al., 2012). Sansalvamide exhibits *in vitro* cytotoxicity against cancer cell lines (Belofsky et al., 1999; Romans-Fuertes et al., 2016), while fusarubin is responsible for perithecial pigmentation of *F. fujikoroi* (Study et al., 2012). Most SB1 chromosomes had at least 66 secreted proteins, but Chr13, Chr14, and Chr15 only had 21, 20, and 40 secreted proteins, respectively (Supplementary Table 1; Figure 2B). For CAZymes, Chr13, Chr14, and Chr15 had three, seven, and six, respectively, compared to other chromosomes with at least 17 (Supplementary Table 1; Figure 2B). The CAZymes identified in these three chromosomes include chitinase 1 (Accession No.: RSL48174.1), chitinase 4 (Accession No.: KAJ4176553.1), glycoside hydrolases (Accession Nos.: XP_046123512.1, XP_046127030.1, XP_046127088.12, KIM92707.1, KAH7017800.1, KAH7009017.1), glycosyl hydrolases (Accession nos.: XP_046125026, XP_046127088.1, XP_046124973.1), glucanase (Accession No.: XP_053015740.1), and pectin lyase (Accession No.: XP_046125011.1). There are at least 21 effectors present in each chromosome except for Chr13 and Chr14 with six (Supplementary Table 1; Figure 2B). Most effectors found in Chr13, Chr14, and Chr15 were uncharacterized or hypothetical except for three out of six from Chr13 and seven out of 21 from Chr15. These effector proteins mainly code for enzymes such as lipase (Accession No.: KAI8649777.1), killer toxin Kp4/SMK (Accession No.: XP_046128691.1), lysophospholipase (Accession No.: XP_018242567.1), Chloroperoxidase (Accession No.: XP_046127231.1), Alpha/Beta hydrolase protein (Accession No.: XP_046127211.1), glycosyl hydrolase family 61-domain-containing protein (Accession No.: XP_046127088.1), pectin lyase fold/virulence factor (Accession No.: XP_046125011.1), tannase and feruloyl esterase-domain-containing protein (Accession No.: XP_046124962.1), and peptidase A4 family-domain-containing protein (Accession No.: XP_046123576.1). We highlight that even with its smallest size and high repeat sequences, Chr15 still has the highest number of genes and more secreted proteins compared to Chr13 and Chr14.

**Figure 2.**
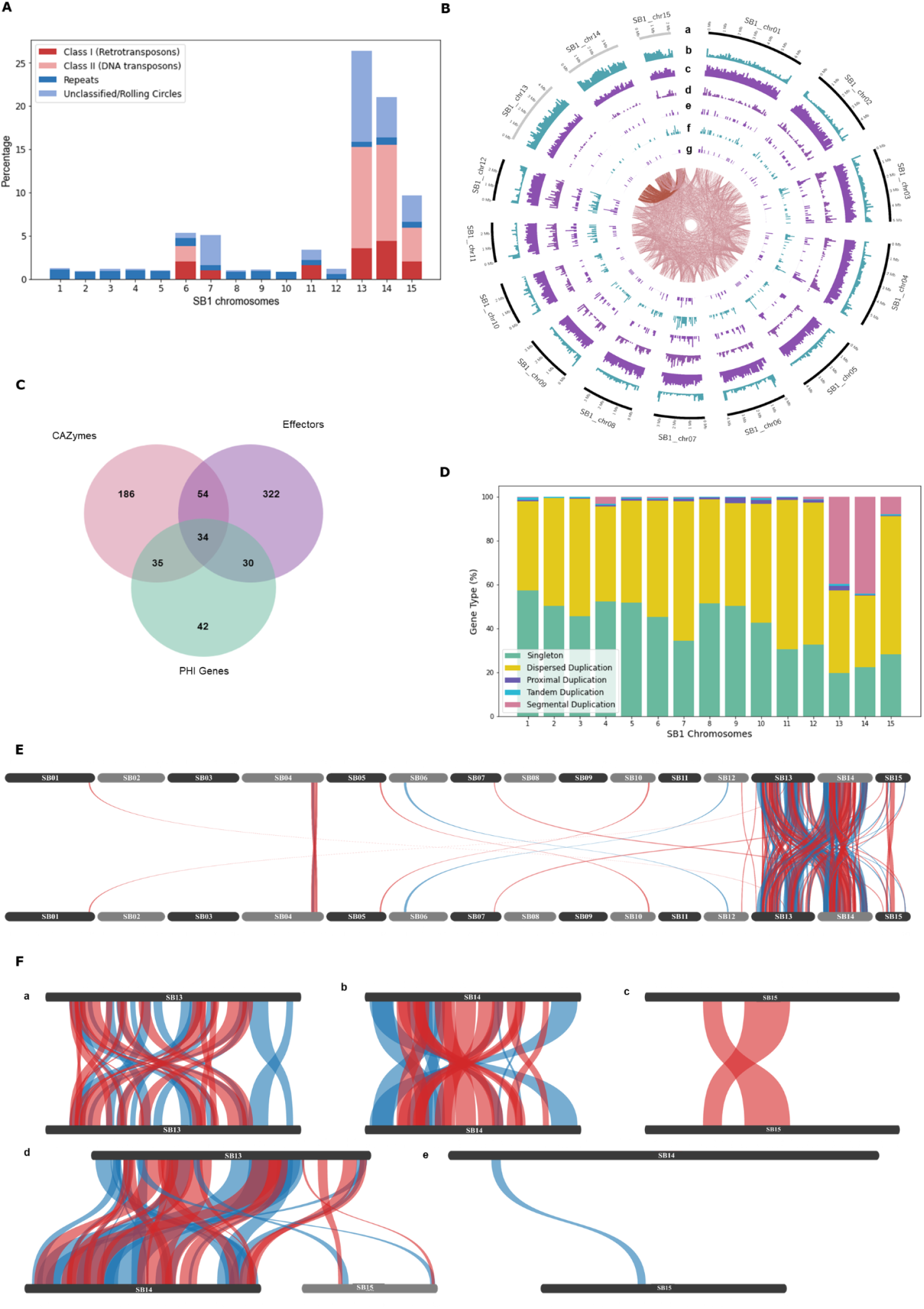
Genomic features of the *Fusarium solani* SB1. **(A)** Repeats and transposable elements of SB1 chromosomes showing Chr13, Chr14, and Chr15 with the highest content among chromosomes; **(B)** Circos plot showing *F. solani* SB1 15 chromosomes at different sizes, gray bands are potential accessory chromosomes (a), repeat element density (b), gene density (c), secreted proteins (d), carbohydrate-active enzymes (e), effectors (f), pathogenicity-host interaction genes (g), and links between SB1 chromosomes. Note the high repeats, fewer genes, CAZymes, effectors, and phi genes in Chr13, Chr14, and Chr15; **(C)** Interaction of CAZymes, effectors, and PHI genes; **(D)** Type of genes in each chromosome of SB1 strain. Note the high percentage of dispersed duplications all over the SB1 genome and segmental duplications within and between Chr13 and Chr14, followed by Chr15 and then Chr04. **(E)** Collinearity of *F. solani* SB1 genome showing duplicated regions, particularly between Chr13, Chr14, and Chr15; **(F)** Close-up view of duplications within and between Chr13, Chr14, and Chr15; Blue and red lines in **E** and **F** are syntenic and inverted sequences, respectively.

Around 69 PHI genes were associated with reduced virulence, five with effector functions, one with loss of pathogenicity, 47 with no effect on pathogenicity, and 19 had mixed functions. Further, around 44% (62) of these proteins are homologous to *Fusarium* species, mainly to *F. graminearum* (36), *F. oxysporum* (12), *F. solani* (12), *F. verticillioides* (1), and *F. virguliforme* (1). The remaining 56% (79) are homologous to proteins found in *Magnaporthe oryzae* (22), *Botrytis sp*. (10), *Colletotrichum sp*. (9), *Trichoderma virens* (9), *Penicillium digitatum* (6), and other organisms (23). A complete list is available in Supplementary Table 2. Focusing on *F. solani* protein homologs, four genes namely PELA (PHI: 179), PELD (PHI: 180), CutA (PHI: 2849), and CSN1 (PHI: 2403), comprise all 12 hits. PELA (PHI:179) and PELD (PHI:180) are essential genes coding for pectate lyase associated with root rots in *Pisum sativum* (Rogers et al., 2000). CutA (PHI:2849) coding for cutinase and CSN1 (PHI: 2403) coding for chitosanase were considered non-essential genes associated with storage rot in *Maxima cucurbita* and *Maxima moscato* (Crowhurst et al., 1997) and root, seedling, and pod rot in *Pisum sativum* (Liu et al., 2010), respectively. No PHIs from Chr13, Chr14, and Chr15 belong to *F. solani*. The number of PHI genes varies across chromosomes, as low as three for Chr15 and as high as 19 for Chr07. Four of 19 PHIs from Chr07 have similarities to *Fusarium solani* essential genes PELA (1) and PELD (3). One particular protein (Chr07_2628) has 99.07% identity with PELD coding for family 3 polysaccharide lyase in *F. solani* (Accession no.: XP_046134888.1). There were no hits of PEP genes in the SB1 secreted proteins.

Considering secreted proteins can be CAZymes, effectors, or PHIs, we determined how many genes have intersecting function. We found 34 genes with all three functions (Figure 2C). Around 80% (27 of 34) of genes code for enzymes such as glycoside hydrolase, pectin/pectate lyase, polygalacturonase, endoglucanase, glucanase, lyases, and cutinase. The remaining genes code for hypothetical and starch-binding domain-containing proteins. Even though most genes are coding for similar functions, they do not have the exact accession numbers in BLAST results and are showing 95-100% query cover and similarity. This displays the arsenal of enzymes encoded in the genome of *F. solani* SB1.

### 3.4 Segmental duplications between and within Chr13 and Chr14

We performed self-alignment of the *F. solani* SB1 DNA sequence and uncovered intriguing inter-chromosomal connections indicative of sequence similarity (Figure 2B, links in the innermost circle). Furthermore, our analysis highlights Chr13’s predominant alignment with Chr14 and Chr15. We further conducted syntenic gene analysis within the *F. solani* SB1 genome, revealing a total of 10,406 dispersed, 195 proximal, 123 tandem, and 963 segmental duplications, alongside 9,101 singletons (Figure 2D). The dispersed duplications explain the links indicating sequence similarity all over the SB1 genome. Chr13 accounts for almost half of the segmental duplications at 44%, followed by Chr14 at 38.5% (Figure 2E). Chr04 also contains segmental duplications at 7%, while Chr15 obtained 5%. The rest of the chromosomes have no segmental duplications. Moreover, our investigation revealed that duplications were predominantly concentrated within and between Chr13 and Chr14 (Figure 2F-a, c, and d), with fewer occurrences observed within Chr15 and between Chr15 and either Chr13 or Chr14 (Figure 2F-c, d, and e).

### 3.5 Comparative Genomics of 12 *Fusarium solani* genomes

#### 3.5.1 Genomic characteristics of *F. solani* group

The genome size of *Fusarium solani* isolates averaged 53.91 Mb, with the smallest at 45.81 Mb from JS-169 and the biggest at 66.64 Mb from CR12 (Figure 3A and Supplementary Table 3). The GC content (%) ranged from 49.5 to 51.5. Genomes achieved 95.6 to 99.7% completeness by BUSCO version 5.2.2 (Simão et al., 2015; Manni et al., 2021). Repeat contents contributed to large sizes of SB1 and CR12 with 11.50 and 17.45%, respectively (Figure 3A). Gene models ranged from 14,663 to 18,410, with an average gene density of 319 ranging from 276 to 334 per Mb where the two largest genomes and highest repeats, CR12 and SB1, also had the lowest gene densities with 276 and 303 per Mb, respectively. Average protein length per genome ranges from 452 to 496 and averages 477 amino acids (AA) (Figure 3B). The protein length distribution warrants good-quality annotation supplementing BUSCO analysis (Nevers et al., 2023).

**Figure 3.**
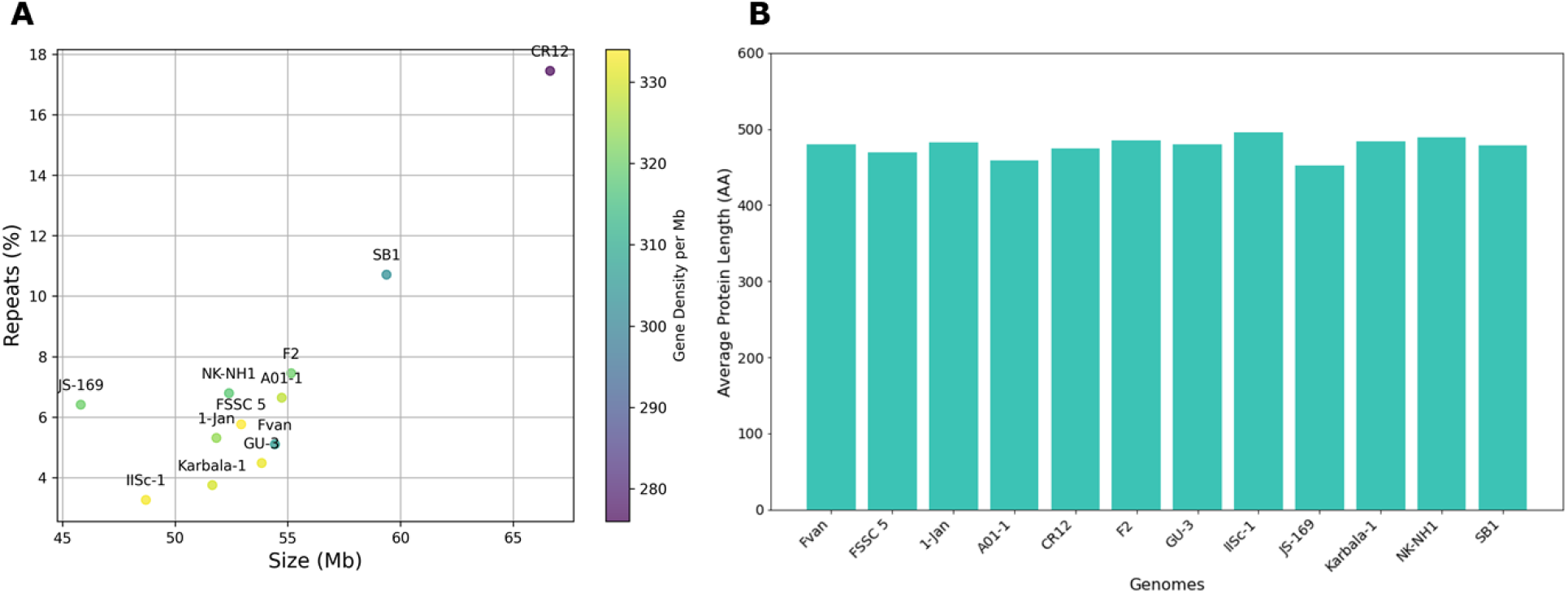
Genomic features of *F. solani* genomes. **(A)** Size (Mb), repeats (%), and gene density (color gradient). The largest genomes, CR12 and SB1, also have the highest repeats but the lowest gene densities; **(B)** Protein length of *F. solani* genomes, averages 477 AA.

#### 3.5.2 Nearly saturated *F. solani* pan-genome

We constructed the pan-genome curve by clustering 204,225 predicted proteins derived from 12 accessions showing decreasing core proteins as new genomes are added. In contrast, new clusters increase with new genomes, indicating an open pan-genome of *Fusarium solani* (Figure 4A). However, we also observed a steady decline of new protein clusters with only under 300 as the 12th genome is added (Figure 4B). This suggests discovering fewer new genes when sequencing additional genomes of *F. solani*. The number of core proteins (present in all strains) is almost the same for all genomes at 11,400 on average, but this varies in percentage relative to the size of the genomes. For example, JS-169 has the lowest number of core proteins (n=11,141) but is equivalent to 76%, the highest among genomes (Figure 4C; Supplementary Table 3). Accessory proteins (present in two or more strains) ranged from 3,217 for JS-169 to 6,417 for SB1, accounting for 21% and 36% of their respective genomes. Unique proteins (present in a single strain) are as low as 97 or 0.6% of F2’s genome and as high as 582 or 3.2% of AO1-1 genomes. The ratio between the number of genes in orthogroups and the number of orthogroups present in a genome is >1 for all strains, indicating that each orthogroup contains more than one gene from each genome (Supplementary File 5). This is shown as paralogs primarily observed in AO1-1 and CR12 at 20% (Figure 4D).

**Figure 4.**
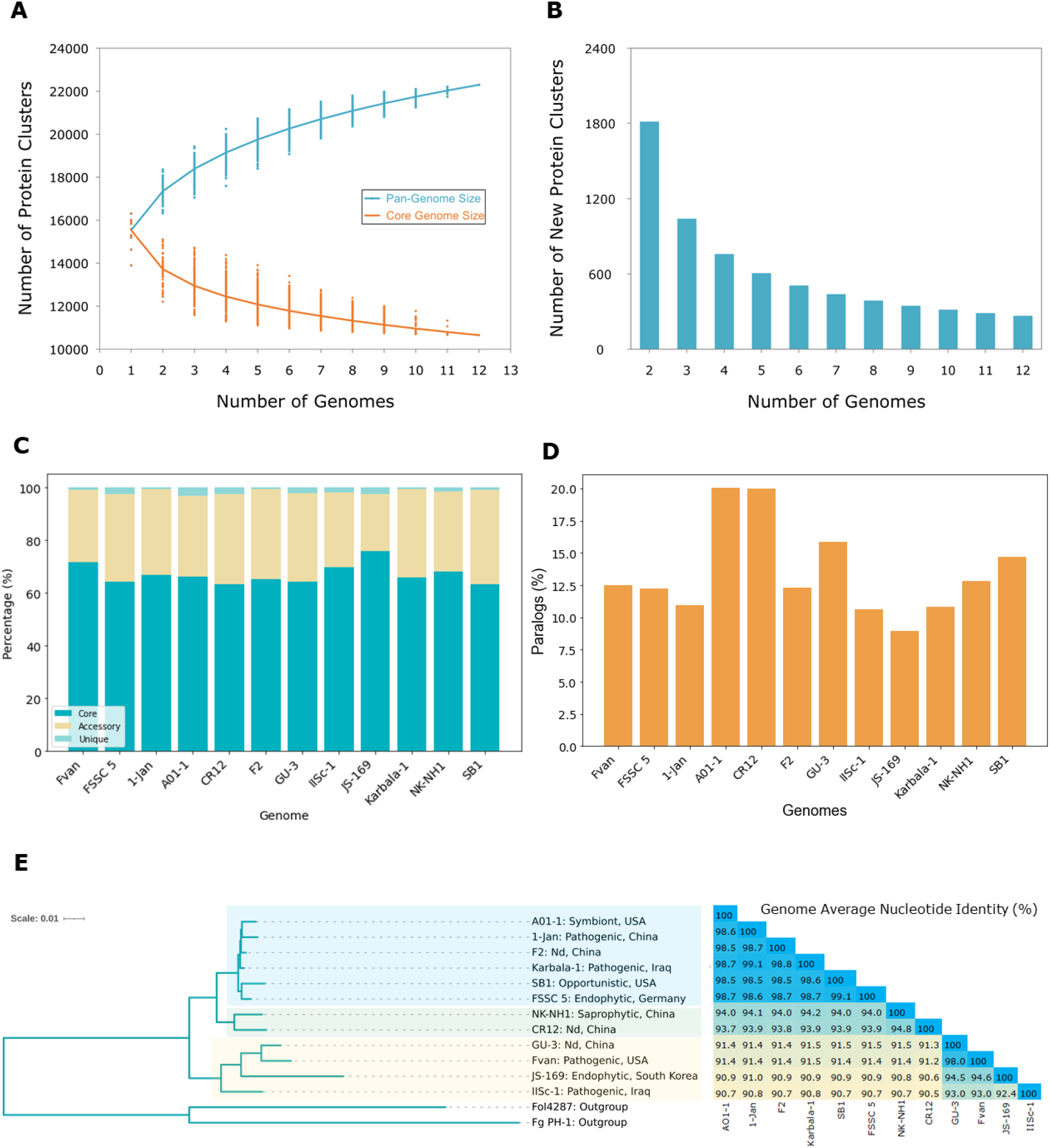
The pan-genome and phylogenetic relationship of *Fusarium solani*. **(A)** The pan-genome and core genome of 12 *F. solani* strains show decreasing core proteins as new genomes are added; **(B)** Number of new protein clusters of *F. solani* in the pan-genome with <300 new protein clusters after the 12th genome; **(C)** Core (present in all strains), accessory (present in two or more strains), and unique (present in one strain) proteins of 12 *F. solani* genomes accounting for an average of 66.7%, 31.6%, and 1.7% of the total pan-genome; **(D)** Percent paralogs (proteins present in more than one copy in an orthogroup) in each genome are primarily found in AO1-1 and CR12; **(E)** Phylogenetic relationships of *F. solani* based on orthogroups (n=10,650) and genome average nucleotide identity (ANI) show this species is not grouped according to lifestyle or origin. Nd, not determined.

#### 3.5.3 Phylogenomic relationships between *F. solani* isolates

OrthoFinder version 2.5.5 (Emms and Kelly 2017; Emms and Kelly, 2019) generated the species tree by utilizing 10,650 orthogroups containing 136,545 genes of *F. solani* strains (Figure 4E). All *F. solani* strains are rooted in the two outgroups Fol4287 and Fg PH1. *F. solani* strains are divided into different clades but it is evident that the strains are not grouped based on their lifestyle or origin. For example, in one clade, the strains AO1-1, 1-Jan, F2, and Karbala-1 are closely related but have different lifestyles and origins. AO1-1 is a symbiont from sweet oranges in the USA, 1-Jan and Karbala-1 are pathogenic isolates from prawns in China and the cockscomb plant in Iraq (Shehan et al., 2023), respectively. F2 was isolated from the rhizospheric soil of Chinese ginseng with a lifestyle that needs to be clearly defined. These four strains share the same ancestor as SB1, an opportunistic isolate from sugarbeet in the USA, and FSSC 5, an endophytic isolate from *Arabidopsis* in Germany. The SB1 isolate was placed in the same clade as FSSC 5, confirming our previous result using the RPB2 gene (Navasca et al., 2023). Another clade sharing the same ancestor as the six previously mentioned strains is two isolates from China, the saprophyte NK-NH1 and CR12, with undetermined lifestyles. The remaining four strains, GU-3, Fvan, JS-169, and IISc-1, with various lifestyles and origins, comprise another clade. The GU3 strain from China and the pea pathogen Fvan from the USA are closely related and share a common ancestor with the mulberry endophytic JS-169 from South Korea. These three strains are related to the evergreen pathogen ISSc-1 from Iraq. Based on the branch lengths, JS-169 had the most significant genetic variation among *F. solani* genomes. The average nucleotide identity (ANI, %) analysis between *F. solani* genomes aligns with the outcomes of the species tree (Figure 4E). The closely related strains AO1-1, 1-Jan, F2, Karbala-1, SB1, and FSSC 5 exhibited a notable similarity, sharing more than 98.5% identity. CR12 and NK-NH1 displayed 93.7 to 94.8% identity with other strains and with each other. GU-3 and Fvan demonstrated a 91.2 to 91.5% similarity with other strains but a higher percentage at 98.0% with each other. JS-169 had comparatively lower ANI with other strains ranging from 90.6 to 91.0% except for its nearest neighbors GU-3 and Fvan, with 94.5% and 94.6%, respectively. IISc-1 exhibited the lowest ANI among strains with 90.5%, except when compared to strains, it shared a common ancestor with – GU-3, Fvan, and JS-169 – with identities ranging from 92.4 to 93.0%.

#### 3.5.4 Enzymatic enrichment of *F. solani* dispensable genome

The top significant pathways enriched by core and dispensable (accessory and unique) genomes are shown in Figure 5. Over 100 core proteins (bars in dark blue in Figure 5A-C) belong to each of the several pathways in the core genome under biological processes (10/10), cellular components (6/10), and molecular function (4/20) while more than 50 proteins mostly occupy the rest. This contrasts with the dispensable genome, where most pathways are occupied by over 25 but less than 50 proteins (bars in lightest blue in Figure 5D-F). While this is the case, fold-enrichment of several pathways is higher in the dispensable genome than in the core genome. The role played by core proteins is crucial in the growth and development of fungi. This is evident in the enrichment of biological processes such as translation and several biosynthetic pathways, as well as activities occurring in the mitochondrion, ribosome, endoplasmic reticulum, and cellular envelopes. On the contrary, pathways enriched in the dispensable genome are more on metabolic pathways and vitamin biosynthetic processes, including cytoplasmic and ribosomal activities. These are additional measures that the fungi would undertake when necessary. We showed the top 10 pathways for biological processes (Figure 5A and 5D) and cellular components (Figure 5B and 5E) and the top 20 for molecular functions (Figure 5C and 5F) to emphasize the enrichment of enzymes in this category, particularly for dispensable proteins. Apart from structural and electron transport activities, six pathways of the core genome are enriched in enzymatic functions. Apart from transporters, 11 pathways are enriched with enzymatic functions in the dispensable genome, including hydrolases, transferases, oxidoreductases, lyases, ligase, isomerase, and dehydrogenase, with the highest fold enrichment at 9.65. All associations between core and dispensable proteins and GO terms are significant, with all FDR values below the threshold (*p*-value: 0.05).

**Figure 5.**
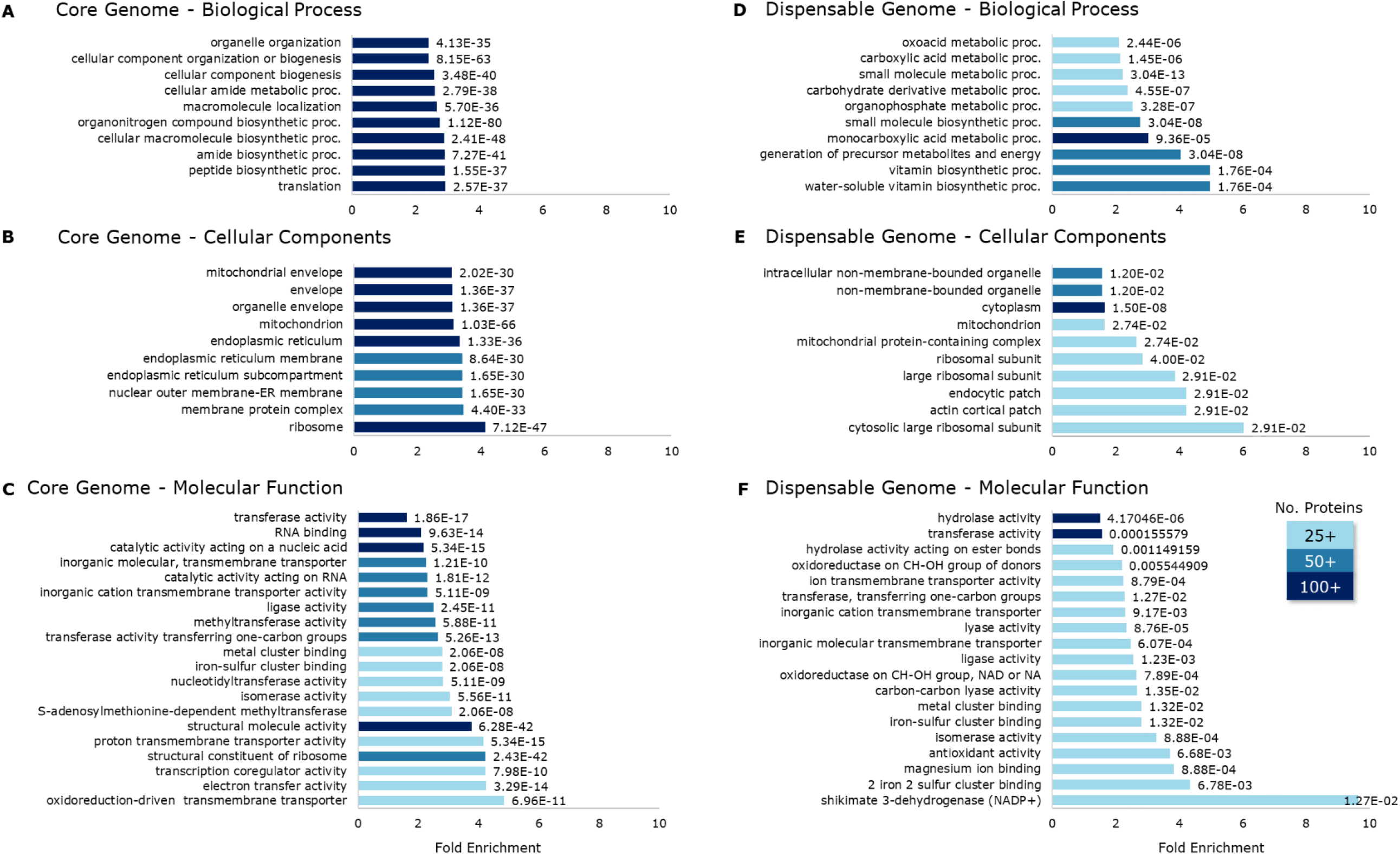
The top gene ontology pathways of core (**A, B, C**) and dispensable (**D, E, F**) genomes are categorized under biological processes (**A, D**), cellular components (**B, E**), and molecular functions (**C, F**). The numbers after the bars represent statistical significance against the FDR threshold (*p*-value: 0.05). The dispensable genome has relatively higher enrichment in several pathways than the core genome. The highest enrichment was observed from shikimate 3-dehydrogenase (9.65) under the molecular function category of the dispensable genome.

## 4.0 Discussion

*Fusarium* species are known to acquire genomic regions, allowing them to infect a wide range of hosts. Members of the *Fusarium solani* species complex (FSSC) are pathogens, saprophytes, and opportunists associated with more than 100 hosts, including plants, animals, and even humans (Coleman et al., 2009). The ability of this group to adapt to different environments reflects their genetic diversity, making them notorious fungal pathogens. Coleman et al. (2009) reported that the expanded genome of *Nectria haematococca* MPVI, now called *F. vanettenii* strain 77-14-3, is due to the specific genes not found in other fungi and single-copy genes occurring in multiple copies. In this study, we identified repeat contents, transposable elements (TEs), and segmental duplications as contributing factors to the expanded genome of the *F. solani* SB1 isolate, particularly by the three chromosomes not found in the reference genome *F. vanettenii* strain 77-13-4. In addition to small size and low GC content, the high TEs and segmental duplications are characteristics of accessory chromosomes (Han et al., 2001; Hatta et al., 2002; Garmaroodi & Taga, 2007; Mehrabi et al., 2011; Coleman et al., 2009; Ma et al., 2010; Yang et al., 2020). A recent study established that segmental duplications shape accessory regions in *F. oxysporum* and *F. solani* (van Westerhoven et al., 2024). With these characteristics, we hypothesize that these three chromosomes, Chr13, Chr14, and Chr15 of the SB1 genome, are accessory chromosomes.

Accessory chromosomes (ACs) are linked to pathogenicity in *F. oxysporum* (Ma et al., 2010; Ma et al., 2023; Yang et al., 2020), but this has not been proven yet in *Nectria haematococca* (syn. *F. solani*). The pea pathogenicity (PEP) and pisatin demethylating ability (PDA) genes are present in the ACs of *F. vanettenii* 77-14-3 (Miao et al., 1991; Kistler et al., 1996; Wasmann and VanEtten, 1996; Han et al., 2001; Liu et al., 2003) and are known to enhance virulence. However, their absence does not significantly impact pathogenicity (Wasmann and VanEtten, 1996; Temporini and VanEtten, 2002). Because we did not find these genes in the SB1 genome, we suspect some other genes harbored by Chr13, Chr14, and Chr15 are responsible for the opportunistic habit of the SB1 isolate. Most genes carried by these chromosomes encode for chitinases 1 and 4, glycoside and glycosyl hydrolases, glucanase, lyase, tannase and esterase, and peptidase. These enzymes are involved in the assembly and degradation of carbohydrates (Lombard et al., 2013) and play a crucial role in pathogenesis by degrading the first line of plant defense, the cell walls (Kubicek et al., 2014). Glycoside hydrolases, an essential enzyme in cell wall degradation, are particularly enhanced in the SB1 genome, potentially aiding the entry of *F. solani* SB1 into sugarbeet.

Segmental duplications are long DNA segments (> 1 Kbp) with high sequence similarity (∼ 90%) along multiple locations in chromosomes (Bailey et al., 2001; Hartmann et al., 2022). They are major sources of evolution and are found in genomes of primates and humans (Bailey et al., 2001; Vollger et al., 2022), *Saccharomyces* species (Dujon et al., 2004), and *Candida albicans* (Rustchenko et al., 1997). In the SB1 genome, massive segmental duplications are found in Chr13 and Chr14. This explains why the collinearity of *F. vanettenii* 77-14-3 Chr14 genes was found on both chromosomes. Because of this large duplication, we deliberated whether Chr13 and Chr14 are identical chromosomes. A closer inspection of the Hi-C contact map (scaffold 5 for Chr13 and scaffold 8 for Chr14), the presence of telomeres, and the genomic characteristics (size, GC content, repeat families, genes) demonstrated they are distinct chromosomes. There are limited explanations for this phenomenon. In *Cryptococcus neoformans*, Fraser et al. (2005) reported a meiotic event where two chromosomes fused, eventually splitting to form two new chromosomes sharing large segmental duplications. Another possible explanation is the involvement of transposable element activity in segmental duplications, as exhibited in the *Fusarium* banana pathogen Tropical Race 4 strain II (van Westerhoven et al., 2024). This is probable since Chr13 and Chr14 have high transposable elements, although we did not determine their proximity to the segmental duplications.

*Fusarium solani* has an open pan-genome but is nearing saturation, with only a few genes uncovered in 12 genomes. We speculate that despite the variation in size observed among *F. solani* genomes, the pan-genome is approaching saturation because the genome expansion is due to the duplication of genes, particularly those coding for enzymes, rather than the emergence of novel genes. This finding is also supported by the enrichment of enzymes in the dispensable genome rather than the core genome. Evident with the strains used in this study, *F. solani* has a wide host range and a diverse lifestyle - pathogens, saprophytes, and opportunistic. The enrichment of several enzymes, such as hydrolases, transferases, oxidoreductases, lyases, ligase, isomerase, and dehydrogenase, apparent by the GO terms in the dispensable genome, supports the ability of *Fusarium solani* to adapt to its varying environment. These enzymes carry out processes essential for adaptations and possibly the reason for the highly adaptive nature of this group.

Most evolutionary relationships are generated using single-copy orthologs, but recent papers argue that some information is lost by not including paralogs (Smith and Hahn, 2021; Smith et al., 2022; Ufimov et al., 2022). This is particularly important when inferring relationships in the concept of adaptation, which is one of the objectives of our study. Moreover, using single-copy orthologs in the phylogenomic analysis might bias the wide range of *F. solani* genomes from 45.81 to 66.64 Mb where paralogs occupy 8-10%. OrthoFinder version 2.5.5 (Emms and Kelly 2017; Emms and Kelly, 2019) generated the species tree using orthogroups (n=10,650) present in all strains of *F. solani*. Around 136,545 genes from these core orthogroups support that *F. solani* is not classified according to lifestyle or origin. The ANI values between *F. solani* genomes corroborate this idea. Our findings supplement those of Hoh et al. (2022), where members of the *Fusarium solani* species complex are not grouped by its animal or plant hosts.

Horizontal gene or chromosome transfer and hybridization between plant pathogenic fungi, especially *Fusarium* species, are ways for pathogens to broaden their host range (Ma et al., 2010; Ma et al., 2013; Mehrabi et al., 2011; Yang et al., 2022). Genome comparisons of other *Fusarium* species, particularly *F. solani*, residing in the soil where other crops, such as potato, dry bean, and soybean, are grown might shed more light on gene and chromosomal transfers within this group. Here, we identified Chr13, Chr14, and Chr15 of the *F. solani* SB1 isolate as potentially accessory chromosomes. Further investigation is needed to discern whether individual or all three chromosomes are necessary for the opportunistic habit of this isolate. The results we presented in this study provide additional evidence of the genome plasticity of the highly adaptive *Fusarium solani*.

## Supporting information

Fusarium solani Pangenome Supplementary Files

## Data Availability Statement

All data relevant to the study are included in the article or uploaded as supplementary information.

## Author Contributions

AN - Investigation, Conceptualization, Methodology, Formal Analysis, Visualization, Validation, Writing - original draft, Writing - reviewing and editing

JS - Methodology, Validation, Writing - reviewing and editing

VS - Methodology, Writing - reviewing and editing

UG - Methodology, Validation, Writing - reviewing and editing

GS - Supervision, Writing - reviewing and editing

TB - Conceptualization, Supervision, Validation, Writing - reviewing and editing

## Acknowledgments

This work used resources of the Center for Computationally Assisted Science and Technology (CCAST) at North Dakota State University, which were made possible in part by NSF MRI Award No. 2019077.

## Supplementary Material

https://docs.google.com/spreadsheets/d/18bMRgZKFpnKLcb1P_xNKrR92hwuWcxf2K1pyspsVIKo/edit#gid=342285514

