## Supplementary material for "Dispensable genome and segmental duplications drive the genome plasticity in *Fusarium solani*": Fusarium solani Pangenome Supplementary Files

**Supplementary Table 1. Genomic features of 15 Fusarium solani strain SB1 chromosomes. Chromosomes in bold are not found in the reference genome F. vanettenii strain 77-13-4.**

| Chr No. | Size (Mb) | GC (%) | Total Repeat Families (%)* | Number of Telomeric Repeats* | Predictions and Annotations* | | | | | | |
|---|---|---|---|---|---|---|---|---|---|---|---|
| | | | | | Protein-coding Genes | Average Gene Density per Mb | Secondary Metabolites | Secreted Proteins | CAZymes | Effectors | PHI |
| Chr01 | 6.54 | 52.55 | 1.23 | 29 | 2,039 | 312 | 6 | 96 | 26 | 35 | 15 |
| Chr02 | 4.69 | 51.74 | 0.93 | 23 | 1,516 | 323 | 4 | 78 | 22 | 23 | 8 |
| Chr03 | 5.1 | 50.53 | 1.17 | 28 | 1,574 | 309 | 6 | 113 | 36 | 32 | 7 |
| Chr04 | 5.89 | 52.05 | 1.21 | 27 | 1,804 | 306 | 2 | 98 | 28 | 37 | 13 |
| Chr05 | 4.12 | 51.84 | 0.97 | 0 | 1,343 | 326 | 1 | 68 | 21 | 28 | 10 |
| Chr06 | 4.03 | 49.59 | 5.3 | 25 | 1,275 | 316 | 5 | 66 | 23 | 25 | 8 |
| Chr07 | 3.39 | 48.39 | 5.04 | 26 | 1,174 | 346 | 6 | 144 | 35 | 62 | 19 |
| Chr08 | 3.49 | 51.23 | 1.02 | 33 | 1,144 | 328 | 2 | 82 | 17 | 28 | 8 |
| Chr09 | 3.24 | 51.09 | 1.11 | 25 | 1,056 | 326 | 0 | 72 | 19 | 22 | 10 |
| Chr10 | 2.94 | 50.65 | 0.84 | 0 | 973 | 331 | 1 | 66 | 13 | 27 | 4 |
| Chr11 | 2.74 | 47.71 | 3.38 | 23 | 914 | 334 | 4 | 93 | 28 | 38 | 9 |
| Chr12 | 2.89 | 48.34 | 1.19 | 28 | 1,033 | 357 | 3 | 120 | 24 | 50 | 18 |
| **Chr13** | **4.44** | **50.60** | **26.34** | **27** | **913** | **206** | **0** | **21** | **3** | **6** | **4** |
| **Chr14** | **3.73** | **50.32** | **21.00** | **26** | **719** | **193** | **0** | **20** | **7** | **6** | **5** |
| **Chr15** | **2.11** | **48.82** | **9.66** | **28** | **512** | **243** | **1** | **40** | **6** | **21** | **3** |
| Total | 59.34 | / | / | / | 17,989 | 303 | 41 | 1,177 | 309 | 440 | 141 |

*Details are available in the Methodology section.

| Supplementary Table 2. PHI Genes hits of F. solani SB1 | | | | | | | | | | | | | | | | |
|---|---|---|---|---|---|---|---|---|---|---|---|---|---|---|---|---|
| # Fields: query id | subject id | | | | | | % identity | alignment length | mismatches | gap opens | q. start | q. end | s. start | s. end | evalue | bit score |
| SB1_chr01_pep_ | Q96TN6 | PHI:256 | GAS1 | 318829 | Magnaporthe_or | reduced_virulen | 62 | 200 | 74 | 2 | 1 | 199 | 20 | 218 | 3.00E-83 | 253 |
| SB1_chr01_pep_ | I1RH33 | PHI:5901 | FGSG_03072 | 5518 | Fusarium_gramir | unaffected_path | 80.24 | 587 | 99 | 6 | 1 | 577 | 21 | 600 | 0 | 971 |
| SB1_chr01_pep_ | Q6IVV6 | PHI:547 | CEL5A | 40559 | Botrytis_cinerea | unaffected_path | 53.75 | 307 | 141 | 1 | 3 | 308 | 118 | 424 | 3.00E-122 | 359 |
| SB1_chr01_pep_ | T2C7K6 | PHI:3226 | Pnl1 | 36651 | Penicillium_digit | reduced_virulen | 58.39 | 310 | 124 | 4 | 5 | 311 | 25 | 332 | 9.00E-117 | 345 |
| SB1_chr01_pep_ | Q5XTQ4 | PHI:541 | LIP1 | 40559 | Botrytis_cinerea | unaffected_path | 58.41 | 553 | 223 | 6 | 7 | 554 | 19 | 569 | 0 | 650 |
| SB1_chr01_pep_ | Q00298 | PHI:69 | CUTA | 40559 | Botrytis_cinerea | unaffected_path | 50.31 | 159 | 78 | 1 | 15 | 172 | 30 | 188 | 9.00E-47 | 154 |
| SB1_chr01_pep_ | A7EB00 | PHI:3936 | Ss-caf1 | 5180 | Sclerotinia_scler | loss_of_pathoge | 56.83 | 183 | 78 | 1 | 1 | 182 | 21 | 203 | 2.00E-71 | 217 |
| SB1_chr01_pep_ | A0A194VQ92 | PHI:10219 | FAEA1_(VM1G_0 | 105487 | Valsa_mali | reduced_virulen | 57.07 | 382 | 137 | 7 | 5 | 373 | 40 | 407 | 4.00E-135 | 395 |
| SB1_chr01_pep_ | G4ZRT3 | PHI:10659 | PsGH7a | 67593 | Phytophthora_sc | reduced_virulen | 51.13 | 442 | 206 | 6 | 1 | 434 | 25 | 464 | 2.00E-154 | 450 |
| SB1_chr01_pep_ | G4MRS8 | PHI:816 | MGG_04582 | 318829 | Magnaporthe_or | reduced_virulen | 50 | 268 | 131 | 2 | 323 | 588 | 273 | 539 | 4.00E-91 | 293 |
| SB1_chr01_pep_ | I1RB96 | PHI:5886 | FGSG_00806 | 5518 | Fusarium_gramir | unaffected_path | 79.12 | 388 | 79 | 2 | 2 | 389 | 28 | 413 | 0 | 581 |
| SB1_chr01_pep_ | A0A1C3YLW9 | PHI:5892 | FGSG_09382 | 5518 | Fusarium_gramir | unaffected_path | 72.37 | 427 | 110 | 3 | 1 | 420 | 21 | 446 | 0 | 601 |
| SB1_chr01_pep_ | Q96VZ3 | PHI:181 | PGX1 | 5507 | Fusarium_oxyspo | unaffected_path | 52.67 | 412 | 189 | 3 | 4 | 409 | 30 | 441 | 1.00E-147 | 430 |
| SB1_chr01_pep_ | I1S1C5 | PHI:5893 | FGSG_10525 | 5518 | Fusarium_gramir | unaffected_path | 62.01 | 379 | 140 | 2 | 10 | 387 | 22 | 397 | 5.00E-166 | 473 |
| SB1_chr01_pep_ | Q8NJ73 | PHI:2043 | XYL-6 | 318829 | Magnaporthe_or | unaffected_path | 65.62 | 32 | 11 | 0 | 393 | 424 | 22 | 53 | 3.00E-09 | 57 |
| SB1_chr07_pep_ | Q96VZ3 | PHI:181 | PGX1 | 5507 | Fusarium_oxyspo | unaffected_path | 56.31 | 412 | 176 | 2 | 11 | 418 | 32 | 443 | 3.00E-165 | 475 |
| SB1_chr07_pep_ | Q04701 | PHI:179 | PELA | 169388 | Fusarium_solani | reduced_virulen | 52.5 | 200 | 91 | 4 | 22 | 218 | 23 | 221 | 3.00E-63 | 199 |
| SB1_chr07_pep_ | Q0R411 | PHI:6417 | Sm1 | 29875 | Trichoderma_vire | reduced_virulen | 73.95 | 119 | 31 | 0 | 2 | 120 | 20 | 138 | 2.00E-60 | 184 |
| SB1_chr07_pep_ | A0A1B5KZM5 | PHI:10357__PHI: | UvHrip1_(UVI_02 | 1159556 | Ustilaginoidea_v | effector_(plant_a | 70.23 | 131 | 39 | 0 | 3 | 133 | 20 | 150 | 3.00E-68 | 205 |
| SB1_chr07_pep_ | O93976 | PHI:2364 | Tom1 | 5507 | Fusarium_oxyspo | reduced_virulen | 74.12 | 313 | 80 | 1 | 7 | 319 | 21 | 332 | 5.00E-173 | 486 |
| SB1_chr07_pep_ | A0A194W2D2 | PHI:10223 | FAEC3_(VM1G_0 | 105487 | Valsa_mali | unaffected_path | 51.6 | 469 | 218 | 3 | 3 | 467 | 35 | 498 | 2.00E-167 | 509 |
| SB1_chr07_pep_ | Q2VLJ1 | PHI:716 | ZEB1 | 5518 | Fusarium_gramir | unaffected_path | 52.1 | 547 | 262 | 0 | 2 | 548 | 17 | 563 | 0 | 623 |
| SB1_chr07_pep_ | Q5KAG0 | PHI:4197 | LHC1 | 5207 | Cryptococcus_ne | reduced_virulen | 62.37 | 372 | 140 | 0 | 4 | 375 | 101 | 472 | 3.00E-170 | 487 |
| SB1_chr07_pep_ | I1RU98 | PHI:6126__PHI:5 | FGSG_07783__S | 5518 | Fusarium_gramir | unaffected_path | 79.18 | 437 | 86 | 2 | 1 | 432 | 22 | 458 | 0 | 733 |
| SB1_chr07_pep_ | Q00845 | PHI:180 | PELD | 169388 | Fusarium_solani | reduced_virulen | 99.07 | 215 | 2 | 0 | 1 | 215 | 19 | 233 | 4.00E-157 | 437 |
| SB1_chr07_pep_ | Q7ZA48 | PHI:323 | VFGLU1 | 93591 | Lecanicillium_fur | reduced_virulen | 61.22 | 410 | 151 | 6 | 1 | 410 | 17 | 418 | 0 | 519 |
| SB1_chr07_pep_ | I1RXJ5 | PHI:9242 | Fghyd3_(FGSG_0 | 5518 | Fusarium_gramir | reduced_virulen | 53.97 | 63 | 28 | 1 | 27 | 89 | 21 | 82 | 1.00E-17 | 71.2 |
| SB1_chr07_pep_ | I1S1J0 | PHI:5894 | FGSG_10595 | 5518 | Fusarium_gramir | unaffected_path | 58.47 | 366 | 147 | 2 | 14 | 379 | 25 | 385 | 1.00E-149 | 431 |
| SB1_chr07_pep_ | G4N8Y3 | PHI:3213 | MoCDIP1 | 318829 | Magnaporthe_or | effector_(plant_a | 70.15 | 335 | 98 | 2 | 2 | 335 | 22 | 355 | 2.00E-159 | 453 |
| SB1_chr07_pep_ | Q00845 | PHI:180 | PELD | 169388 | Fusarium_solani | reduced_virulen | 55.19 | 212 | 80 | 6 | 7 | 215 | 26 | 225 | 5.00E-67 | 207 |
| SB1_chr07_pep_ | Q00845 | PHI:180 | PELD | 169388 | Fusarium_solani | reduced_virulen | 50.47 | 212 | 95 | 4 | 21 | 227 | 27 | 233 | 7.00E-58 | 189 |
| SB1_chr07_pep_ | G4N3U4 | PHI:5188 | MoHPX1 | 318829 | Magnaporthe_or | reduced_virulen | 52.58 | 213 | 94 | 4 | 4 | 212 | 33 | 242 | 2.00E-64 | 201 |
| SB1_chr07_pep_ | Q2VLJ1 | PHI:716 | ZEB1 | 5518 | Fusarium_gramir | unaffected_path | 60.29 | 544 | 212 | 1 | 5 | 548 | 24 | 563 | 0 | 702 |
| SB1_chr07_pep_ | G4N4F9 | PHI:2166__PHI:6 | CBP1 | 318829 | Magnaporthe_or | unaffected_path | 55.32 | 47 | 18 | 2 | 224 | 269 | 406 | 450 | 2.00E-06 | 47 |
| SB1_chr05_pep_ | I1S2L3 | PHI:7283 | Pg1 | 5518 | Fusarium_gramir | reduced_virulen | 91.04 | 335 | 30 | 0 | 1 | 335 | 25 | 359 | 0 | 614 |
| SB1_chr05_pep_ | Q8NJS2 | PHI:697 | Ugt51E1 | 5022 | Leptosphaeria_m | unaffected_path | 70.98 | 379 | 107 | 2 | 2 | 378 | 19 | 396 | 0 | 579 |
| SB1_chr05_pep_ | I1RX03 | PHI:3985 | FGSG_08844_(Sc | 5518 | Fusarium_gramir | unaffected_path | 50.85 | 59 | 18 | 1 | 20 | 78 | 454 | 501 | 6.00E-07 | 49.7 |
| SB1_chr05_pep_ | O59928 | PHI:144 | CHT42 | 29875 | Trichoderma_vire | reduced_virulen | 64.13 | 407 | 139 | 2 | 2 | 401 | 24 | 430 | 0 | 555 |
| SB1_chr05_pep_ | Q04701 | PHI:179 | PELA | 169388 | Fusarium_solani | reduced_virulen | 69.57 | 230 | 70 | 0 | 150 | 379 | 13 | 242 | 3.00E-119 | 348 |
| SB1_chr05_pep_ | G4MKI0 | PHI:2118__PHI:5 | MgSM1__MSP1 | 318829 | Magnaporthe_or | effector_(plant_a | 60.83 | 120 | 46 | 1 | 1 | 120 | 19 | 137 | 2.00E-49 | 156 |
| SB1_chr05_pep_ | I1S0V3 | PHI:5902 | FGSG_10343 | 5518 | Fusarium_gramir | unaffected_path | 53.27 | 612 | 240 | 6 | 2 | 589 | 23 | 612 | 0 | 664 |
| SB1_chr05_pep_ | A9QUB2 | PHI:2403 | CSN1 | 169388 | Fusarium_solani | increased_virule | 50.41 | 121 | 60 | 0 | 3 | 123 | 22 | 142 | 5.00E-40 | 136 |
| SB1_chr05_pep_ | G2X4G0 | PHI:11606 | Vd424Y | 27337 | Verticillium_dahl | effector_(plant_a | 79.79 | 188 | 38 | 0 | 1 | 188 | 36 | 223 | 6.00E-107 | 308 |
| SB1_chr05_pep_ | I1R9N9 | PHI:5881 | PRB1 | 5518 | Fusarium_gramir | reduced_virulen | 86.38 | 514 | 64 | 3 | 6 | 514 | 21 | 533 | 0 | 914 |
| SB1_chr12_pep_ | X0J8P9 | PHI:5393 | GLX | 5507 | Fusarium_oxyspo | reduced_virulen | 77.94 | 902 | 179 | 3 | 1 | 902 | 21 | 902 | 0 | 1492 |
| SB1_chr12_pep_ | A0A098DJU9 | PHI:9062__PHI:1 | KatG2_(FGSG_12 | 5518 | Fusarium_gramir | reduced_virulen | 80.42 | 756 | 147 | 1 | 3 | 758 | 20 | 774 | 0 | 1253 |
| SB1_chr12_pep_ | K9G4Z7 | PHI:10923 | PDIG_23520_(Sc | 36651 | Penicillium_digit | unaffected_path | 65.52 | 116 | 37 | 1 | 5 | 120 | 35 | 147 | 4.00E-52 | 163 |
| SB1_chr12_pep_ | Q96VZ3 | PHI:181 | PGX1 | 5507 | Fusarium_oxyspo | unaffected_path | 78.48 | 446 | 78 | 2 | 5 | 450 | 28 | 455 | 0 | 714 |
| SB1_chr12_pep_ | Q00845 | PHI:180 | PELD | 169388 | Fusarium_solani | reduced_virulen | 77.67 | 215 | 46 | 1 | 1 | 213 | 19 | 233 | 3.00E-124 | 353 |
| SB1_chr12_pep_ | Q5KAG0 | PHI:4197 | LHC1 | 5207 | Cryptococcus_ne | reduced_virulen | 59.89 | 374 | 148 | 1 | 8 | 381 | 101 | 472 | 2.00E-164 | 472 |
| SB1_chr12_pep_ | I1S368 | PHI:6127 | PLC | 5518 | Fusarium_gramir | unaffected_path | 80.77 | 598 | 115 | 0 | 3 | 600 | 50 | 647 | 0 | 1055 |
| SB1_chr12_pep_ | I1RU06 | PHI:6124__PHI:5 | FGSG_07678__S | 5518 | Fusarium_gramir | reduced_virulen | 83.58 | 408 | 67 | 0 | 5 | 412 | 17 | 424 | 0 | 693 |
| SB1_chr12_pep_ | Q0E7H5 | PHI:2341 | BeNEP1 | 278938 | Botrytis_elliptica | unaffected_path | 58.45 | 219 | 88 | 1 | 10 | 225 | 27 | 245 | 1.00E-94 | 279 |
| SB1_chr12_pep_ | Q6IVV6 | PHI:547 | CEL5A | 40559 | Botrytis_cinerea | unaffected_path | 53.59 | 306 | 137 | 4 | 68 | 368 | 118 | 423 | 1.00E-113 | 340 |

| | | | | | | | | | | | | | | | | |
|---|---|---|---|---|---|---|---|---|---|---|---|---|---|---|---|---|
| SB1_chr12_pep_ | K9G4Z7 | PHI:10923 | PDIG_23520_(Sc | 36651 | Penicillium_digit | unaffected_path | 70.83 | 120 | 34 | 1 | 10 | 128 | 28 | 147 | 8.00E-58 | 178 |
| SB1_chr12_pep_ | I1RHV1 | PHI:6123__PHI:5 | FGSG_03366__S | 5518 | Fusarium_gramir | reduced_virulen | 74.79 | 468 | 113 | 2 | 6 | 473 | 23 | 485 | 0 | 766 |
| SB1_chr12_pep_ | G4N8Y3 | PHI:3213 | MoCDIP1 | 318829 | Magnaporthe_or | effector_(plant_a | 75.22 | 335 | 83 | 0 | 1 | 335 | 21 | 355 | 7.00E-177 | 497 |
| SB1_chr12_pep_ | I1RHI8 | PHI:6121 | FGSG_03243 | 5518 | Fusarium_gramir | reduced_virulen | 67.92 | 533 | 167 | 3 | 2 | 533 | 21 | 550 | 0 | 762 |
| SB1_chr12_pep_ | Q9C4A0 | PHI:2817 | Aapg1 | 5599 | Alternaria_altern | unaffected_path | 68.19 | 349 | 107 | 3 | 11 | 357 | 32 | 378 | 4.00E-156 | 446 |
| SB1_chr12_pep_ | I1RDQ0 | PHI:9241 | Fghyd2_(FGSG_0 | 5518 | Fusarium_gramir | reduced_virulen | 52.17 | 115 | 47 | 2 | 3 | 110 | 24 | 137 | 2.00E-34 | 117 |
| SB1_chr12_pep_ | Q6WIH5 | PHI:4970 | MEP1 | 523103 | Trichophyton_m | reduced_virulen | 57.12 | 625 | 252 | 6 | 3 | 622 | 19 | 632 | 0 | 738 |
| SB1_chr12_pep_ | G8AA67 | PHI:2476 | CcpelA | 27358 | Colletotrichum_c | reduced_virulen | 61.25 | 271 | 101 | 3 | 16 | 283 | 56 | 325 | 3.00E-112 | 329 |
| SB1_chr09_pep_ | E3QFA7 | PHI:10749 | CLU5d_(GLRG_0 | 31870 | Colletotrichum_g | reduced_virulen | 53.11 | 177 | 76 | 3 | 19 | 194 | 65 | 235 | 1.00E-52 | 177 |
| SB1_chr09_pep_ | A0A0C4DHY4 | PHI:4880 | Pgx6 | 5507 | Fusarium_oxyspo | unaffected_path | 82.86 | 420 | 72 | 0 | 1 | 420 | 21 | 440 | 0 | 754 |
| SB1_chr09_pep_ | G4MXR2 | PHI:8753__PHI:8 | MoChia1_(MGG_ | 318829 | Magnaporthe_or | reduced_virulen | 58.86 | 367 | 146 | 3 | 8 | 370 | 29 | 394 | 3.00E-157 | 450 |
| SB1_chr09_pep_ | J4W023 | PHI:7181 | CypB | 176275 | Beauveria_bassia | reduced_virulen | 84.44 | 180 | 28 | 0 | 1 | 180 | 29 | 208 | 4.00E-112 | 320 |
| SB1_chr09_pep_ | Q00845 | PHI:180 | PELD | 169388 | Fusarium_solani | reduced_virulen | 63.18 | 220 | 71 | 5 | 1 | 215 | 19 | 233 | 5.00E-88 | 261 |
| SB1_chr09_pep_ | G4ML12 | PHI:6739 | MoGls2 | 318829 | Magnaporthe_or | reduced_virulen | 66.18 | 958 | 296 | 7 | 1 | 933 | 25 | 979 | 0 | 1342 |
| SB1_chr09_pep_ | A0A384JSB2 | PHI:10365 | BcBGL1_(BCIN_0 | 40559 | Botrytis_cinerea | reduced_virulen | 58.75 | 766 | 300 | 10 | 31 | 781 | 33 | 797 | 0 | 872 |
| SB1_chr09_pep_ | Q6WER3 | PHI:432__PHI:42 | FGL1__Fgl1 | 5518 | Fusarium_gramir | reduced_virulen | 66.27 | 332 | 109 | 3 | 1 | 332 | 16 | 344 | 7.00E-162 | 459 |
| SB1_chr09_pep_ | X0MKY5 | PHI:11176 | FoEG1_(FOTG_02 | 5507 | Fusarium_oxyspo | unaffected_path | 69.09 | 220 | 68 | 0 | 2 | 221 | 30 | 249 | 5.00E-109 | 316 |
| SB1_chr09_pep_ | Q8NJ73 | PHI:2043 | XYL-6 | 318829 | Magnaporthe_or | unaffected_path | 62.5 | 32 | 12 | 0 | 442 | 473 | 22 | 53 | 1.00E-07 | 52 |
| SB1_chr11_pep_ | I1RIQ3 | PHI:1575 | GzOB015 | 5518 | Fusarium_gramir | unaffected_path | 76.13 | 222 | 52 | 1 | 1 | 222 | 21 | 241 | 7.00E-123 | 351 |
| SB1_chr11_pep_ | I1RJ45 | PHI:6120 | FGSG_03846 | 5518 | Fusarium_gramir | reduced_virulen | 70.12 | 425 | 104 | 2 | 3 | 404 | 22 | 446 | 0 | 621 |
| SB1_chr11_pep_ | A0A0C4DHY4 | PHI:4880 | Pgx6 | 5507 | Fusarium_oxyspo | unaffected_path | 62.53 | 443 | 152 | 5 | 1 | 442 | 21 | 450 | 0 | 588 |
| SB1_chr11_pep_ | I1RHQ4 | PHI:5898 | FGSG_03315 | 5518 | Fusarium_gramir | unaffected_path | 60.2 | 402 | 148 | 1 | 2 | 403 | 22 | 411 | 7.00E-173 | 492 |
| SB1_chr11_pep_ | X0MKY5 | PHI:11176 | FoEG1_(FOTG_02 | 5507 | Fusarium_oxyspo | unaffected_path | 69.82 | 222 | 65 | 2 | 1 | 221 | 29 | 249 | 3.00E-108 | 314 |
| SB1_chr11_pep_ | K9G4Z7 | PHI:10923 | PDIG_23520_(Sc | 36651 | Penicillium_digit | unaffected_path | 65.12 | 129 | 44 | 1 | 1 | 128 | 19 | 147 | 5.00E-56 | 173 |
| SB1_chr11_pep_ | G4ZRT3 | PHI:10659 | PsGH7a | 67593 | Phytophthora_sc | reduced_virulen | 52.04 | 442 | 202 | 6 | 1 | 434 | 25 | 464 | 3.00E-161 | 473 |
| SB1_chr11_pep_ | Q04701 | PHI:179 | PELA | 169388 | Fusarium_solani | reduced_virulen | 95.56 | 225 | 10 | 0 | 1 | 225 | 18 | 242 | 3.00E-159 | 443 |
| SB1_chr11_pep_ | I1RGU8 | PHI:5897 | FGSG_02976 | 5518 | Fusarium_gramir | unaffected_path | 82.42 | 364 | 64 | 0 | 2 | 365 | 26 | 389 | 0 | 615 |
| SB1_chr06_pep_ | Q99174 | PHI:2849 | CutA | 169388 | Fusarium_solani | unaffected_path | 98.51 | 201 | 3 | 0 | 1 | 201 | 30 | 230 | 2.00E-143 | 402 |
| SB1_chr06_pep_ | G4MVX4 | PHI:3216 | MoCDIP4 | 318829 | Magnaporthe_or | effector_(plant_a | 70.59 | 34 | 10 | 0 | 284 | 317 | 262 | 295 | 4.00E-09 | 55.5 |
| SB1_chr06_pep_ | Q9C1F9 | PHI:566 | Cel2 | 5017 | Bipolaris_zeicola | unaffected_path | 53.37 | 401 | 176 | 8 | 1 | 397 | 18 | 411 | 2.00E-143 | 418 |
| SB1_chr06_pep_ | Q5XTQ4 | PHI:541 | LIP1 | 40559 | Botrytis_cinerea | unaffected_path | 52.94 | 544 | 248 | 5 | 1 | 542 | 31 | 568 | 0 | 564 |
| SB1_chr06_pep_ | G4NIS9 | PHI:785 | MGG_04128 | 318829 | Magnaporthe_or | reduced_virulen | 70.7 | 529 | 146 | 5 | 3 | 523 | 23 | 550 | 0 | 761 |
| SB1_chr06_pep_ | A9QUB2 | PHI:2403 | CSN1 | 169388 | Fusarium_solani | increased_virule | 96.81 | 282 | 8 | 1 | 1 | 282 | 20 | 300 | 0 | 557 |
| SB1_chr06_pep_ | I1S5M1 | PHI:5903 | FGSG_12142 | 5518 | Fusarium_gramir | unaffected_path | 71.06 | 577 | 165 | 2 | 1 | 576 | 19 | 594 | 0 | 858 |
| SB1_chr06_pep_ | I1RFN8 | PHI:1166 | ATG15 | 5518 | Fusarium_gramir | reduced_virulen | 81.73 | 591 | 103 | 2 | 1 | 586 | 46 | 636 | 0 | 938 |
| SB1_chr13_pep_ | O59928 | PHI:144 | CHT42 | 29875 | Trichoderma_vir | reduced_virulen | 59.78 | 363 | 138 | 2 | 1 | 357 | 23 | 383 | 2.00E-161 | 462 |
| SB1_chr13_pep_ | Q6WER3 | PHI:432__PHI:42 | FGL1__Fgl1 | 5518 | Fusarium_gramir | reduced_virulen | 56.1 | 328 | 129 | 3 | 1 | 314 | 16 | 342 | 1.00E-131 | 381 |
| SB1_chr13_pep_ | O59928 | PHI:144 | CHT42 | 29875 | Trichoderma_vir | reduced_virulen | 55.37 | 363 | 128 | 5 | 1 | 331 | 23 | 383 | 5.00E-140 | 407 |
| SB1_chr13_pep_ | O59928 | PHI:144 | CHT42 | 29875 | Trichoderma_vir | reduced_virulen | 59.51 | 410 | 158 | 2 | 1 | 404 | 23 | 430 | 0 | 515 |
| SB1_chr02_pep_ | E3QFA7 | PHI:10749 | CLU5d_(GLRG_0 | 31870 | Colletotrichum_g | reduced_virulen | 50.75 | 67 | 31 | 1 | 405 | 469 | 407 | 473 | 6.00E-14 | 72.8 |
| SB1_chr02_pep_ | Q9P470 | PHI:257 | GAS2 | 318829 | Magnaporthe_or | reduced_virulen | 54.78 | 272 | 106 | 2 | 1 | 272 | 22 | 276 | 1.00E-88 | 268 |
| SB1_chr02_pep_ | G3F417 | PHI:3703 | Fvtox1 | 232082 | Fusarium_virguli | reduced_virulen | 91.5 | 153 | 13 | 0 | 1 | 153 | 20 | 172 | 5.00E-103 | 295 |
| SB1_chr02_pep_ | W7M3S4 | PHI:7144 | FvSCP1 | 117187 | Fusarium_vertici | reduced_virulen | 63.95 | 319 | 101 | 5 | 1 | 305 | 18 | 336 | 1.00E-131 | 380 |
| SB1_chr02_pep_ | I1RYH0 | PHI:6409 | FgCdc11 | 5518 | Fusarium_gramir | reduced_virulen | 97.33 | 375 | 10 | 0 | 66 | 440 | 6 | 380 | 0 | 748 |
| SB1_chr02_pep_ | V9MGK6 | PHI:4096 | FoOCH1 | 5507 | Fusarium_oxyspo | reduced_virulen | 86.21 | 319 | 43 | 1 | 1 | 319 | 42 | 359 | 0 | 577 |
| SB1_chr02_pep_ | O59939 | PHI:222 | PELB | 474922 | Colletotrichum_g | reduced_virulen | 66.29 | 264 | 86 | 2 | 33 | 295 | 68 | 329 | 1.00E-119 | 349 |
| SB1_chr02_pep_ | G4NA54 | PHI:2214 | Endo-1_4-beta-x | 318829 | Magnaporthe_or | reduced_virulen | 68.45 | 187 | 59 | 0 | 2 | 188 | 43 | 229 | 1.00E-92 | 277 |
| SB1_chr14_pep_ | O59928 | PHI:144 | CHT42 | 29875 | Trichoderma_vir | reduced_virulen | 59.51 | 410 | 158 | 2 | 1 | 404 | 23 | 430 | 0 | 515 |
| SB1_chr14_pep_ | O59928 | PHI:144 | CHT42 | 29875 | Trichoderma_vir | reduced_virulen | 59.76 | 410 | 157 | 2 | 1 | 404 | 23 | 430 | 0 | 517 |
| SB1_chr14_pep_ | Q9C1F9 | PHI:566 | Cel2 | 5017 | Bipolaris_zeicola | unaffected_path | 53.87 | 401 | 174 | 8 | 1 | 397 | 18 | 411 | 3.00E-145 | 423 |
| SB1_chr14_pep_ | O59928 | PHI:144 | CHT42 | 29875 | Trichoderma_vir | reduced_virulen | 56.02 | 407 | 174 | 4 | 4 | 407 | 25 | 429 | 1.00E-167 | 479 |
| SB1_chr14_pep_ | O59928 | PHI:144 | CHT42 | 29875 | Trichoderma_vir | reduced_virulen | 58.13 | 363 | 143 | 3 | 1 | 356 | 23 | 383 | 8.00E-154 | 442 |
| SB1_chr08_pep_ | I1RXJ5 | PHI:9242 | Fghyd3_(FGSG_0 | 5518 | Fusarium_gramir | reduced_virulen | 64.41 | 59 | 21 | 0 | 3 | 61 | 24 | 82 | 5.00E-21 | 79 |
| SB1_chr08_pep_ | G4MQW5 | PHI:7320 | GH18 | 318829 | Magnaporthe_or | unaffected_path | 57.25 | 393 | 154 | 7 | 10 | 391 | 44 | 433 | 8.00E-158 | 454 |
| SB1_chr08_pep_ | G4NCF7 | PHI:2138 | SGA1 | 318829 | Magnaporthe_or | unaffected_path | 53.27 | 107 | 46 | 2 | 252 | 354 | 542 | 648 | 9.00E-27 | 110 |
| SB1_chr08_pep_ | G4NGA7 | PHI:2032 | VTL1 | 318829 | Magnaporthe_or | unaffected_path | 53.82 | 498 | 226 | 2 | 20 | 516 | 42 | 536 | 0 | 573 |

| | | | | | | | | | | | | | | | | |
|---|---|---|---|---|---|---|---|---|---|---|---|---|---|---|---|---|
| SB1_chr08_pep_ | W6ZBM1 | PHI:4989 | Ppt1 | 101162 | Bipolaris_oryzae | reduced_virulen | 50.71 | 282 | 129 | 4 | 8 | 279 | 28 | 309 | 8.00E-101 | 301 |
| SB1_chr08_pep_ | Q8J0H8 | PHI:7228 | Tre1 | 318829 | Magnaporthe_or | unaffected_path | 66.12 | 670 | 220 | 3 | 1 | 664 | 23 | 691 | 0 | 945 |
| SB1_chr08_pep_ | Q04701 | PHI:179 | PELA | 169388 | Fusarium_solani | reduced_virulen | 67.43 | 218 | 68 | 2 | 1 | 218 | 28 | 242 | 1.00E-102 | 299 |
| SB1_chr08_pep_ | K9G4Z7 | PHI:10923 | PDIG_23520_(Sc | 36651 | Penicillium_digit | unaffected_path | 62.02 | 129 | 48 | 1 | 1 | 129 | 20 | 147 | 1.00E-51 | 162 |
| SB1_chr10_pep_ | A7F946 | PHI:2411 | Ss-ggt1 | 5180 | Sclerotinia_scler | reduced_virulen | 57.53 | 365 | 145 | 3 | 205 | 560 | 130 | 493 | 2.00E-138 | 413 |
| SB1_chr10_pep_ | A0A0N9DQW5 | PHI:10255 | VmXyl1 | 105487 | Valsa_mali | reduced_virulen | 57.26 | 365 | 141 | 3 | 1 | 365 | 20 | 369 | 4.00E-139 | 414 |
| SB1_chr10_pep_ | I1RDW3 | PHI:9245 | Fghyd5_(FGSG_0 | 5518 | Fusarium_gramir | reduced_virulen | 85.14 | 74 | 11 | 0 | 1 | 74 | 25 | 98 | 7.00E-45 | 141 |
| SB1_chr10_pep_ | I1RHQ4 | PHI:5898 | FGSG_03315 | 5518 | Fusarium_gramir | unaffected_path | 76.94 | 386 | 87 | 1 | 8 | 393 | 25 | 408 | 0 | 600 |
| SB1_chr04_pep_ | A0A194W2D2 | PHI:10223 | FAEC3_(VM1G_0 | 105487 | Valsa_mali | unaffected_path | 61.49 | 470 | 175 | 4 | 1 | 465 | 31 | 499 | 0 | 607 |
| SB1_chr04_pep_ | A0A2H3U592 | PHI:11257 | FoMep1 | 5507 | Fusarium_oxyspo | reduced_virulen | 73.25 | 243 | 63 | 2 | 4 | 246 | 35 | 275 | 2.00E-135 | 385 |
| SB1_chr04_pep_ | I1RPX8 | PHI:1581 | GzOB021 | 5518 | Fusarium_gramir | unaffected_path | 53.63 | 248 | 114 | 1 | 1 | 248 | 27 | 273 | 3.00E-93 | 281 |
| SB1_chr04_pep_ | E3QHX9 | PHI:6227 | KRE5 | 31870 | Colletotrichum_g | reduced_virulen | 64.29 | 1473 | 473 | 8 | 2 | 1426 | 25 | 1492 | 0 | 1968 |
| SB1_chr04_pep_ | A0A0D2XVZ5 | PHI:5236 | Dnj1 | 5507 | Fusarium_oxyspo | reduced_virulen | 80 | 500 | 94 | 3 | 1 | 494 | 22 | 521 | 0 | 827 |
| SB1_chr04_pep_ | G2WWK9 | PHI:2712 | NLP2 | 27337 | Verticillium_dahl | reduced_virulen | 67.45 | 212 | 67 | 1 | 1 | 210 | 28 | 239 | 1.00E-95 | 281 |
| SB1_chr04_pep_ | O60038 | PHI:1034 | Cpcat1 | 5111 | Claviceps_purpu | unaffected_path | 76.21 | 681 | 158 | 3 | 2 | 681 | 37 | 714 | 0 | 1088 |
| SB1_chr04_pep_ | I1RR94 | PHI:6122__PHI:5 | FGSG_06610__S | 5518 | Fusarium_gramir | reduced_virulen | 77 | 626 | 116 | 2 | 2 | 627 | 21 | 618 | 0 | 995 |
| SB1_chr04_pep_ | A0A384JWC5 | PHI:10367 | BcBGL3_(BCIN_1 | 40559 | Botrytis_cinerea | reduced_virulen | 65.92 | 851 | 270 | 9 | 19 | 864 | 56 | 891 | 0 | 1154 |
| SB1_chr04_pep_ | I1RR60 | PHI:5899 | FGSG_06572 | 5518 | Fusarium_gramir | unaffected_path | 72.65 | 852 | 233 | 0 | 14 | 865 | 29 | 880 | 0 | 1320 |
| SB1_chr04_pep_ | L2FHG9 | PHI:3972 | CDA | 474922 | Colletotrichum_g | unaffected_path | 64.46 | 363 | 122 | 2 | 12 | 367 | 22 | 384 | 3.00E-175 | 499 |
| SB1_chr04_pep_ | G4MTF8 | PHI:2204 | Endo-1_4-beta-x | 318829 | Magnaporthe_or | reduced_virulen | 68.98 | 303 | 94 | 0 | 1 | 303 | 29 | 331 | 1.00E-149 | 426 |
| SB1_chr04_pep_ | Q0E7H5 | PHI:2341 | BeNEP1 | 278938 | Botrytis_elliptica | unaffected_path | 59.17 | 218 | 86 | 1 | 1 | 218 | 32 | 246 | 2.00E-91 | 271 |
| SB1_chr03_pep_ | Q6WER3 | PHI:432__PHI:42 | FGL1__Fgl1 | 5518 | Fusarium_gramir | reduced_virulen | 56.79 | 324 | 126 | 2 | 5 | 314 | 19 | 342 | 2.00E-131 | 381 |
| SB1_chr03_pep_ | G8AA67 | PHI:2476 | CcpelA | 27358 | Colletotrichum_c | reduced_virulen | 77.63 | 295 | 66 | 0 | 1 | 295 | 31 | 325 | 3.00E-168 | 473 |
| SB1_chr03_pep_ | L7JCL9 | PHI:10459 | Gtb1 | 318829 | Magnaporthe_or | reduced_virulen | 62.38 | 529 | 195 | 3 | 2 | 529 | 23 | 548 | 0 | 681 |
| SB1_chr03_pep_ | G4N4F9 | PHI:2166__PHI:6 | CBP1 | 318829 | Magnaporthe_or | unaffected_path | 61.11 | 36 | 14 | 0 | 32 | 67 | 415 | 450 | 7.00E-08 | 51.6 |
| SB1_chr03_pep_ | I1RW12 | PHI:5890 | FGSG_08464 | 5518 | Fusarium_gramir | unaffected_path | 60.37 | 429 | 164 | 2 | 10 | 432 | 20 | 448 | 2.00E-180 | 514 |
| SB1_chr03_pep_ | E3QFA6 | PHI:10751 | CLU5c_(GLRG_04 | 31870 | Colletotrichum_g | unaffected_path | 65.81 | 272 | 93 | 0 | 1 | 272 | 20 | 291 | 2.00E-133 | 382 |
| SB1_chr03_pep_ | E3QFA7 | PHI:10749 | CLU5d_(GLRG_04 | 31870 | Colletotrichum_g | reduced_virulen | 65.87 | 208 | 69 | 1 | 63 | 270 | 67 | 272 | 2.00E-88 | 281 |
| SB1_chr15_pep_ | T2C7K6 | PHI:3226 | Pnl1 | 36651 | Penicillium_digit | reduced_virulen | 58.79 | 347 | 139 | 2 | 1 | 347 | 20 | 362 | 3.00E-134 | 390 |
| SB1_chr15_pep_ | Q5XTQ4 | PHI:541 | LIP1 | 40559 | Botrytis_cinerea | unaffected_path | 56.7 | 552 | 231 | 4 | 1 | 547 | 20 | 568 | 0 | 630 |
| SB1_chr15_pep_ | A0A194W2D2 | PHI:10223 | FAEC3_(VM1G_0 | 105487 | Valsa_mali | unaffected_path | 53.5 | 471 | 212 | 5 | 4 | 468 | 28 | 497 | 4.00E-176 | 532 |

| Supplementary Table 3. Information on Fusarium solani strains used in pan-genom | | | | | | | | | | | | | |
|---|---|---|---|---|---|---|---|---|---|---|---|---|---|
| **Strain** | **NCBI Accession No.** | **Size (Mb)** | **N50 (Mb)** | **L50** | **GC%** | **Proteins** | **Gene Density per Mb** | **Repeats** | **Genome Completeness (%, proteins)** | **Source** | **Lifestyle** | **Origin** | **Reference** |
| Fvan 77-13-4 | GCA_000151355 | 54.43 | 1.3 | 10 | 50.8 | 15705 | 307 | 5.1 | 99.3 | Pea (Pisum sativum) | Pathogenic | USA | Unpublished at time of study |
| FSSC 5 MPI-SDFR-AT-0091 | GCA_020744495.1 | 52.93 | 2.2 | 9 | 51 | 17654 | 334 | 5.76 | 99.7 | Healthy Arabidopsis plant | Endophytic | Germany | Mesny et. al, 2021 |
| SB1 | GCA_023522795.1 | 59.38 | 4 | 6 | 50.5 | 17989 | 303 | 10.71 | 98.5 | Galled sugarbeet (Beta vulgaris) | Opportunistic | USA | Navasca et al., 2023; this study |
| NK-NH1 | GCA_033085375.1 | 52.39 | 3.9 | 6 | 51.5 | 16644 | 318 | 6.79 | 99.5 | Hull wood of Nanhai NO.1 Shipwreck | Saprophytic | China | Unpublished at time of study |
| CR12 | GCA_030014125.1 | 66.64 | 3.5 | 8 | 49.5 | 18410 | 276 | 17.45 | 98.9 | Root of Ginkgo biloba | Nd | China | Unpublished at time of study |
| JS-169 | GCA_002215905.1 | 45.81 | 2.6 | 6 | 50 | 14663 | 320 | 6.41 | 95.6 | Twig of Mulberry (Morus alba) | Endophytic | South Korea | Kim et al., 2017 |
| 1-Jan | GCA_019320015.1 | 51.83 | 2 | 9 | 51.5 | 16772 | 324 | 5.31 | 97.1 | Broodstock of Fannabin prawns (Penaeus vannamei) | Pathogenic | China | Unpublished at time of study |
| IISc-1 | GCA_013168735.1 | 48.7 | 1.6 | 9 | 50.5 | 16197 | 333 | 3.26 | 99.5 | Stem cuttings of evergreen (Taxus celebica) | Endophytic | India | Unpublished at time of study |
| Karbala-1 | GCA_024220475.1 | 51.65 | 780.4 Kb | 19 | 51.5 | 17036 | 330 | 3.75 | 99.3 | Cockscomb (Celosia argentea L.) | Pathogenic | Iraq | Shehan et al., 2023 |
| A01-1 | GCA_027945525.1 | 54.73 | 373.3 Kb | 44 | 50.5 | 17956 | 328 | 6.64 | 99.6 | Root/soil of sweet orange (Citrus sinensis) | Symbiont | USA | Unpublished at time of study |
| F2 | GCA_029603225.1 | 55.15 | 357.9 Kb | 47 | 50.5 | 17654 | 320 | 7.45 | 99.2 | Rhizosphere soil of Chinese ginseng (Panax notoginseng) | Nd | China | Unpublished at time of study |
| GU-3 | GCA_027574645.1 | 53.84 | 186.6 Kb | 80 | 51.5 | 17879 | 332 | 4.48 | 99.6 | Chinese liquorice (Glycyrrhiza uralensis) | Nd | China | Unpublished at time of study |

**Supplementary Table 4. Core, Accessory, and Unique Genes of *Fusarium solani* strains**

| Genome | Genes | | |
|---|---|---|---|
| | Core | Accessory | Unique |
| Fvan | 11,254 | 4,311 | 140 |
| Fusso | 11,333 | 5,884 | 437 |
| 1-Jan | 11,202 | 5,471 | 99 |
| A01-1 | 11,910 | 5,464 | 582 |
| CR12 | 11,660 | 6,309 | 441 |
| F2 | 11,317 | 5,906 | 97 |
| GU-3 | 11,475 | 6,012 | 392 |
| IISc-1 | 11,279 | 4,628 | 290 |
| JS-169 | 11,141 | 3,127 | 395 |
| Karbala-1 | 11,236 | 5,671 | 129 |
| NK-NH1 | 11,350 | 5,051 | 243 |
| SB1 | 11,388 | 6,417 | 184 |

**Supplementary Table 5. Orthologous Genes of *Fusarium solani* Pangenome**

| Description | Fvan | FSSC 5 | 1-Jan | A01-1 | CR12 | F2 | GU-3 | IISc-1 | JS-169 | Karbala-1 | NK-NH1 | SB1 |
|---|---|---|---|---|---|---|---|---|---|---|---|---|
| Number of genes | 15,705 | 17,654 | 16,772 | 17,956 | 18,410 | 17,320 | 17,879 | 16,197 | 14,663 | 17,036 | 16,644 | 17,989 |
| Number of genes in orthogroups | 15,600 | 17,240 | 16,686 | 17,429 | 18,055 | 17,227 | 17,574 | 15,912 | 14,281 | 16,948 | 16,435 | 17,858 |
| Number of unassigned genes | 105 | 414 | 86 | 527 | 355 | 93 | 305 | 285 | 382 | 88 | 209 | 131 |
| Percentage of genes in orthogroups | 99.3 | 97.7 | 99.5 | 97.1 | 98.1 | 99.5 | 98.3 | 98.2 | 97.4 | 99.5 | 98.7 | 99.3 |
| Percentage of unassigned genes | 0.7 | 2.3 | 0.5 | 2.9 | 1.9 | 0.5 | 1.7 | 1.8 | 2.6 | 0.5 | 1.3 | 0.7 |
| Number of orthogroups containing species | 14,528 | 15,895 | 15,502 | 15,337 | 15,526 | 15,847 | 15,662 | 14,902 | 13,513 | 15,743 | 15,086 | 15,880 |
| Percentage of orthogroups containing species | 75.2 | 82.3 | 80.3 | 79.4 | 80.4 | 82 | 81.1 | 77.2 | 70 | 81.5 | 78.1 | 82.2 |
| Number of species-specific orthogroups | 11 | 10 | 6 | 23 | 38 | 2 | 38 | 2 | 6 | 17 | 15 | 20 |
| Number of genes in species-specific orthogroups | 35 | 23 | 13 | 55 | 86 | 4 | 87 | 5 | 13 | 41 | 34 | 53 |
| Percentage of genes in species-specific orthogroups | 0.2 | 0.1 | 0.1 | 0.3 | 0.5 | 0 | 0.5 | 0 | 0.1 | 0.2 | 0.2 | 0.3 |
| Ratio (Number of genes in orthogroups / Number of orthogroups containing species) | 1.07 | 1.08 | 1.08 | 1.14 | 1.16 | 1.09 | 1.12 | 1.07 | 1.06 | 1.08 | 1.09 | 1.12 |
